# Prior selection affects phenotypic and transcriptional response to hypoxia

**DOI:** 10.1101/2022.03.13.484120

**Authors:** Millicent N. Ekwudo, Morad C. Malek, Cora E. Anderson, Lev Y. Yampolsky

**Affiliations:** Department of Biological Sciences, East Tennessee State University, Johnson City TN 37614, USA

## Abstract

Hypoxia has profound and diverse effects on aerobic organisms, disrupting oxidative phosphorylation and activating several protective pathways. Predictions have been made that exposure to mild intermittent hypoxia may be protective against more severe exposure and may extend lifespan. Both effects are likely to depend on prior selection on phenotypic and transcriptional plasticity in response to hypoxia, and may therefore show signs of local adaptation. Here we report the lifespan effects of chronic, mild, intermittent hypoxia (CMIH) and short-term survival in acute severe hypoxia (ASH) in four clones of *Daphnia magna* originating from either permanent or intermittent habitats, the latter regularly drying up with frequent hypoxic conditions. We show that CMIH extended the lifespan in the two clones originating from intermittent habitats but had the opposite effect in the two clones from permanent habitats, which also showed lower tolerance to ASH. Exposure to CMIH did not protect against ASH; to the contrary, *Daphnia* from the CMIH treatment had lower ASH tolerance than normoxic controls. Few transcripts changed their abundance in response to the CMIH treatment in any of the clones. After 12 hours of ASH treatment, the transcriptional response was more pronounced, with numerous protein-coding genes with functionality in mitochondrial and respiratory metabolism, oxygen transport, and gluconeogenesis showing up-regulation. While clones from intermittent habitats showed somewhat stronger differential expression in response to ASH than those from permanent habitats, there were no significant hypoxia-by-habitat of origin or CMIH-by-ASH interactions. GO enrichment analysis revealed a possible hypoxia tolerance role by accelerating the molting cycle and regulating neuron survival through up-regulation of cuticular proteins and neurotrophins, respectively.

## Introduction

A vast literature exists on local adaptation to hypoxia in terrestrial habitats where low oxygen partial pressure is a permanent feature of high altitudes (Hochachka et al. 1998; Yi et al. 2010; Scott et al. 2015; Brutsaert et al. 2019). Much less is known about local adaptation and phenotypic plasticity in response to hypoxia in aquatic habitats, where oxygen availability is both lower and more variable than in the atmosphere. Oxygen concentration can vary from levels above the 8-10 mg/L saturation levels during peaks of photosynthetic activity to below 1 mg/L in warm, stalled, and organic-rich habitats.

Oxygen concentrations are also variable in time, both on dial and seasonal scale. Species greatly differ in their ability to survive in hypoxic conditions, but data on intraspecific heritable or phenotypic variation in hypoxia tolerance are fragmentary and largely limited to fish (Gorr et al. 2010; Dammark et al. 2018; Brennan et al. 2018; Andersen et al. 2020; Borowiec & Scott 2020; Crispo et al. 2020) rather than invertebrates (Gorr et al. 2010; Sandoval-Castillo et al. 2018; Falfushynska et al. 2020). Not uncommonly, little evidence of local adaptation is discovered despite existing environmental oxygen availability gradients (Dammark et al. 2018), indicating possible selective constraints. In contrast, data on the acclimation effects of mild or intermittent hypoxia that allow higher tolerance to more severe hypoxic treatments have indicated that at least some such hormesis effects exist (Yang et al. 2013; Hermes-Lima et al. 2015; Borowiec & Scott 2020; Peruzza, 2021). However, details of transcriptional and biochemical plasticity behind these acclimation effects are not well understood.

In general, regulatory responses to hypoxia have been well characterized.

Hypoxia-mediated responses are controlled by a family of heterodimeric transcription factors, the hypoxia-inducible factors (HIFs), which interact with numerous regulatory pathways and are conserved across Metazoa (Yeo 2019). These pathways include SIRT1, AMPK, mTOR, and NF-κ B pathways (Ruderman et al. 2010; Gorr et al. 2010; Hong et al. 2014; Antikainen et al. 2017; Pan et al. 2017). Hypoxic responses at the cellular level are maintained mostly by HIF-1α (Wang et al. 1995; Iyer et al. 1998). HIF-1α is constitutively expressed, although in normoxia the protein is quickly degraded through a proteolysis pathway initiated by the oxygen-sensing action of HIF proline hydroxylase.

Additionally, HIF expression is up-regulated under hypoxia and/or by increased ROS conditions through NF-κB (a redox-sensitive transcription factor; Bonello et al. 2007). HIF-1α in turn, acts as a transcription factor for a variety of hypoxia-responsive genes by binding to Hypoxia Response Elements (HREs) located in their promoter regions. HIF-1α exerts transcriptional control over roughly 100 target genes during hypoxia (Yang et al. 2015), especially genes associated with oxygen transport and homeostasis such as hemoglobins, erythropoietin (*EPO*), and Vascular Endothelial Growth Factor (*VEGF*), that are vital for increasing tissue perfusion and oxygenation as an adaptive response to hypoxia (Zeis et al. 2009; Yeo 2019).

HIF-1α also upregulates lactate dehydrogenase and, consequently, the conversion of pyruvate to lactate when low oxygen hinders membrane phosphorylation, thereby decreasing the NAD^+^/NADH ratio. This ratio is vital for intracellular redox homeostasis, especially in the mitochondria and nucleus, as well as cellular signaling and the regulation of metabolic activities. Importantly, HIF1α also modulates the cell cycle, through c-MYC and IGF2 signaling and apoptotic pathways (Yeo 2019). Furthermore, hypoxia induces inflammation and the immune response by increasing the expression of tumor necrosis factor-alpha (TNF- α (Scholz et al. 2013).

Several interconnected downstream pathways are known to offer protection against oxidative damage and extend lifespan, including FOXO, AMPK, and sirtuin regulatory pathways. Members of the FOXO gene family pathway (including *C. elegans* daf-16) promote longevity by up-regulating DNA repair (Tran et al. 2002) and catalase activity (Wang et al. 2017; Murtaza et al. 2017. The AMP-activated protein kinase (AMPK) pathway regulates oxidative energy metabolism and responds to a variety of stresses such as a decrease in glucose levels, ischemia, heat shock, and hypoxia (Yeo 2019). Hypoxia can directly trigger the AMPK pathway when there is an elevation in the AMP:ATP ratio. Previous studies have implicated this pathway in slowing the rate of aging by decreasing oxidative stress through increased thioredoxin levels and the autophagic degradation of protein aggregates (Salminen et al. 2012). AMPK also plays a role in the regulation of mitochondrial biogenesis through PGC-1α (Jäger et al. 2007) and in cellular autophagy (Gui et al. 2017) via direct phosphorylation of the autophagy-promoting protein kinase ULK1, or indirectly through down-regulating the mTOR pathway (Gorr et al. 2010; Kim et al. 2011; Shang et al. 2011). AMPK also attenuates endoplasmic reticulum (ER) stress and modulates inflammation by suppressing NF- B κ signaling. which in turn inhibits NF- κ -mediated inflammation (Salminen et al. 2011).

This anti-inflammatory activity of AMPK is thought to contribute to healthspan and lifespan extension (Yeo 2019).

Finally, sirtuins, a family of proteins homologous to the silent information regulator 2 (sir2) protein of yeast, are NAD^+^-dependent protein deacetylases (and deacylases) whose activities have attracted attention because of their roles in inflammation, apoptosis, and senescence. Hypoxia is known to up-regulate the sirtuin pathway via HIF-1a activity (Rui et al. 2011), protecting cells against age-associated pathologies by promoting DNA stability, oxidative stress alleviation, and regulation of glucose and lipid metabolism (Guarente 2011; Kilic et al. 2015). Importantly, the sirtuin pathway can enhance mitochondrial biogenesis by deacetylating p53 and PGC-1α (Lee and Gu 2013). The sirtuin pathway is interconnected with FOXO and AMPK pathways, as sirtuins can upregulate both. In turn, the AMPK activity can result in the phosphorylation of sirtuins, p53, PGC-1α and FOXO proteins.

As the data on the complex interplay between these diverse hypoxia-initiated protective pathways has accumulated in the last 2 decades, it has become common to hypothesize that chronic exposure to mild and/or intermittent hypoxia may extend lifespan and hormetically acclimate organisms to tolerating more severe or more prolonged hypoxic conditions. It is not clear, though, whether such responses show significant variation in nature, and if so, whether such variation reflects local adaptation. Data on transcriptional responses to hypoxia has just started to accumulate, through both transcriptome-wide studies (Mu et al. 2020; Tian et al. 2020; Zhou et al. 2020; Feng et al. 2021; Hu et al. 2021; Jie et al. 2021; Kim et al. 2021; Xu L. et al. 2021; Xu Y. et al. 2021) and targeted qPCR-based studies focused on anaerobic metabolism-related genes (Aksakal E, Ekinci D. 2021; Amorim et al. 2021). These studies reveal that signals of some of the hypoxia-inducible pathways described above are detectable in fish and invertebrates and that several downstream pathways showing response include anaerobic glycolysis and immune responses. Because these studies seldom analyze differential expression in different genotypes or geographically distinct populations (but see Zhou et al. 2020; Kim et al. 2021), the question of genetic variation for hypoxia response remains open.

The goal of this study is to detect local adaptation and adaptive plasticity in response to hypoxia on a phenotypic and transcriptional level using the freshwater zooplankton crustacean *Daphnia magna* Straus as a model organism. Cladoceran crustaceans including *Daphnia* reproduce by cyclic parthenogenesis, which makes them ideal models for studying the response to environments, as genetically identical, and yet outbred individuals can be maintained in different experimental conditions. *D. magna* genome is relatively well characterized and orthologs of many genes known to participate in hypoxia response are known. The HIF-1α induction of hemoglobin expression has been well characterized in *Daphnia* (Pirow et al. 2001; Zeis et al. 2003; Gorr et al 2004; Colbourne et al. 2011; Klumpen et al. 2017). A moderately long exposure to hypoxia can increase mortality rate and decrease body size and fecundity (Seidl et al. 2005; Lyu et al. 2015). On the other hand, a brief exposure to mild hypoxia may increase heat tolerance (Coggins et al. 2017), possibly through activation of antioxidant pathways (Klumpen et al. 2017). Previously published data on transcriptional responses to severe chronic hypoxia in *Daphnia* embryos indicated up-regulation of hemoglobin genes, HSPs, glutathione metabolism-related transcripts, and down-regulation of vitellogenins, histones, and histone modification related transcripts (Lai et al. 2016). These effects may be trans-generational (Lai et al. 2016).

Here we report differential long-term survival in chronic mild intermittent hypoxia (a decrease of O2 concentration from 8 to 4 mg/L twice daily throughout the lifespan; hereafter, CMIH treatment) and short-term survival in acute severe hypoxia (1.5 mg/L for 36 hours; hereafter, ASH treatment) in four geographically distinct *D. magna* clones. Two of these clones originated from intermittent water bodies that regularly dry out in summer, with *Daphnia* populations recolonizing them from resting eggs. Periods immediately preceding drying out are likely to be characterized by overheating and hypoxia. The other two clones originated from permanent temperate water bodies that are unlikely to experience hypoxic conditions regularly. Thus, we can compare transcriptional responses to hypoxia in genotypes with different selection histories with respect to hypoxia tolerance. We analyze differential transcription in all four combinations of control vs. hypoxia in chronic vs. acute treatments, along with data on feeding and respiration rate, and lactate/pyruvate ratio, which can be used as a proxy for NADH/NAD+ ratio.

## Materials and Methods

### Clones’ provenance and maintenance

The geographic origins of the four clones used in this study are listed in Supplementary Table S1. Clones were obtained from Basel University (Switzerland) Daphnia Stock Collection (Dieter Ebert, personal communication) and maintained in the lab in COMBO water (Kilham et al. 1998) at 20 °C under a 16:8 D:L light cycle and fed by a diet of *Scenedesmus acutus.* Prior to all experiments, experimental cohorts were established through the following procedure: Five randomly selected grandmother females from each clone were maintained from birth till their 3^rd^ clutch of offspring were born at the density of 1 individual per 20 ml of COMBO water with *Scenedesmus* food added daily to the concentration of 10^5^ cells/ml and water changed every four days.

**Table 1.** Likelihood Ratio proportional hazards tests of the effects of habitat of origin type, clones nested within habitats, hypoxic conditions, and their interactions on Daphnia lifespan (Fig. 1). Top: comparison of CMIH (4 mg O_2_/L twice daily) to normoxic (8 mg O_2_/L) control. Middle: same with the switch from CMIH to a normoxic regime at day 30 included. Bottom: the same analysis was conducted separately for each habitat type. In the analysis with the switch treatment included, the significant effect of hypoxia treatment is due to both the difference between the switch and CMIH treatment (p<0.05) and between the switch and control (p<0.0001).

| Source | DF | Log Ratio $\chi^2$ | P value |
| --- | --- | --- | --- |
| Lifespan in CMIH vs. normoxic control |  |  |  |
| habitat | 1 | 32.11 | <0.0001 |
| clone[habitat] | 2 | 23.78 | <0.0001 |
| CMIH | 1 | 10.97 | 0.0009 |
| habitat*CMIH | 1 | 8.93 | 0.0028 |
| clone*CMIH [habitat] | 2 | 2.19 | 0.33 |
| Lifespan in CMIH vs. normoxic control vs. CMIH->normoxia switch |  |  |  |
| habitat | 1 | 24.44 | <0.0001 |
| clone[habitat] | 2 | 4.45 | 0.1083 |
| CMIH | 2 | 56.38 | <0.0001 |
| habitat*CMIH | 2 | 9.24 | 0.0099 |
| clone*CMIH [habitat] | 4 | 13.39 | 0.0095 |
| Habitat of origin: intermittent |  |  |  |
| clone | 1 | 1.07 | 0.30 |
| CMIH | 2 | 30.87 | <.0001 |
| clone*CMIH | 2 | 8.272 | 0.016 |
| Habitat of origin: permanent |  |  |  |
| clone | 1 | 1.50 | 0.22 |
| CMIH | 2 | 43.10 | <.0001 |
| clone*CMIH | 2 | 4.69 | 0.096 |

Offspring from their 2^nd^ and 3^rd^ clutches were used to establish the maternal generation, which was in turn maintained in the same conditions until enough offspring from the second or consecutive clutches could be collected to form the experimental cohorts. Neonates were maintained in groups of 20 in 200 mL jars with COMBO water for the first 6 days of their lives until transferred to the corresponding experimental tanks.

### Chronic mild intermittent hypoxia treatment

*Daphnia* cohorts were maintained in 5-L tanks each containing eight plastic containers with 1 mm nylon mesh bottoms, which allowed free water exchange and the removal of neonates during water changes. Water volume and daily food ratios were adjusted every 4 days to maintain 20 ml of water per individual, and 10^5^ *Scenedesmus* cells were added per mL per day. Chronic mild intermittent hypoxia treatment (CMIH thereafter) was achieved by bubbling N_2_ through the experimental tanks with continuous monitoring of oxygen concentration by the Extech DO210 probe (Nashua, NH, USA) twice daily until the concentration was lowered to 4 mg/L. At the same time, the control tanks were aerated with ambient air until the oxygen concentration reached 8 mg/L. Between these procedures, the oxygen concentration in the hypoxia tanks typically raised to 6.5 mg/L by diffusion, whereas the control tanks typically experienced a drop in oxygen concentration of 7–7.5 mg/L due to respiration of *Daphnia*. To investigate whether early life exposure to mild hypoxia provides the same protection as life-time exposure, one of the replicates within the CMIH treatment was switched to normoxic treatment on day 30 of the experiment. Because this switch treatment included only one replicate, the results of this comparison will be interpreted as highly speculative and preliminary.

The water was changed, neonates removed, and a census of *Daphnia* cohorts conducted every three days. At various time points, individuals from each cohort were sampled for body size, fecundity, feeding rate and respiration rate measurements (with replacement), or for lactate and pyruvate concentration determination, acute hypoxia tolerance experiment, or RNAseq (without replacement).

The CMIH experiment was conducted twice, with each block consisting of two CMIH and two control tanks (plus one CMIH to control the switch replicate in one of the blocks).

### Lactate and Pyruvate Measurements

Lactate and pyruvate assays were conducted using CellBiolab kits on 15 to 20 day old and 55 to 60 day old *Daphnia* from two clones (GB and IL) sampled from the experimental tanks and stored frozen at -80°C until assay time. Each *Daphnia* was homogenized in 100 µL ice-cold PBS with a pestle, and the homogenates were centrifuged at 4°C. 25 µL of supernatant were pipetted into each of the lactate and pyruvate assay plates using the manufacturer’s protocol (CellBiolab catalog #s 101820174 & 82320181). Additionally, two replicate aliquots of 15 µL of the supernatant each were used to quantify soluble proteins by Bradford assay, with 185 µl of Bradford colorimetric reagent added to each well. All assay well plates were analyzed using a BIOTEK plate reader (Agilent, Santa Clara, CA, USA). Lactate and pyruvate fluorescence assay sensitivity was set to 35, and the Bradford assay absorption was measured at 595 nm.

### Respiration and filtering rates measurements

Respiration rate was measured in individual *Daphnia* sampled from the cohorts by placing them in either 200 μ or 1700 μ 24-well respirometry glass plates (Loligo ®, Denmark), sealing them with PCR sealing tape, and measuring oxygen concentration using SDR fluorescence sensors (PreSens, Germany). Either normoxic (8 mg O_2_/L) or hypoxic (4 mg O_2_/L) COMBO water was used in these measurements. Each measurement continued for 45 – 60 minutes, with oxygen concentration recorded every 15 seconds until the oxygen concentration dropped by at least 1 mg/L. The first 15 minutes after the transfer of *Daphnia* into the respirometry plates were discarded as the break-in period. The slope of the linear regression of oxygen concentration over time was used as the measure of oxygen consumption rate.

Feeding (filtering) rate was measured by placing individual *Daphnia* into plastic 24-well plates with normoxic COMBO water containing the initial concentration of 2x10^5^ *Scenedesmus* cells per mL and measuring chlorophyll fluorescence at the start and after 8, 12, and 24 hours of filtering activity.

After both respiration and feeding rate measurements, *Daphnia* were measured, weighed to determine their wet weight to the precision of 0.1 mg, and returned to the experimental tanks.

### Mitochondrial membrane potential

Samples for mitochondrial membrane potential (ΔΨ_m_) measurements were taken from the cohorts at the median lifespan of all cohorts, i.e., at the age of 80 days to maximize longevity-related effects. ΔΨ_m_ was measured by means of rhodamine-123 assays as described in Anderson et al (2022). Briefly, *Daphnia* were exposed for 24 h to a M solution of rhodamine-123 dye, washed 3 times, and photographed using a Leica μ DM3000 microscope with a 10x objective (0.22 aperture) equipped with a Leica DFc450C color camera, with a 488 nm excitation / broadband (>515 nm) emission filter. The fluorescence from the following tissues and organs was measured: antenna-driving muscle, epipodite, brain, and optical lobe, with the median intensity of fluorescence used as the measure of ΔΨ_m_. We have previously shown that ΔΨ_m_ does not change much with age in most tissues (Anderson et al. 2022), so it is likely that these measurements are representative of any age of *Daphnia* exposed to CMIH for a sufficient time.

### Acute Severe Hypoxia Experiments and Sample Collection for RNA-Seq

For acute severe hypoxia (ASH) tolerance measurement, 25 day old *Daphnia* were sampled from each of the four tanks in one of the two CMIH experiment blocks. They were moved into 70 mL cell culture flasks filled with COMBO water with the concentration of oxygen at or below 1 mg/L, sealed without air bubbles, 7 *Daphnia* per flask, 5 replicate flasks per clone per treatment. Flasks were kept at 20°C. The acute hypoxia experiment was set up at 9:00 p.m. and mortality was recorded 12 hours later and every hour thereafter. Individuals for RNA sequencing were frozen at the beginning of the experiment (controls) and after 12 hours of exposure (before any mortality occurred), sampling 2 individuals from each flask (ASH treatment). Flasks were then topped with 1 mg O_2_/L water and sealed again.

### RNA Sequencing

Two individuals from each of the four clones (IL, FI, GB, and HU) and from each of the hypoxia treatments were frozen during the acute hypoxia experiment (see above). The four treatments were: *Daphnia* reared at normoxia, *Daphnia* reared at normoxia and exposed to acute hypoxia for 12 hours, *Daphnia* reared in CMIH, and *Daphnia* reared in CMIH and exposed to acute hypoxia for 12 hours. RNA was extracted using Qiagen RNAeasy kit (Cat ID: 74134) and quantified using a Qubit (ThermoFisher) fluorometer.

Following extraction, RNAs were reverse transcribed and sequencing libraries were constructed from the cDNAs as prescribed by the Oxford Nanopore Technology (Oxford, UK) PCR-cDNA Barcoding kit protocol (SQK-PCB109), with 3 biological replicates per clone per treatment, each replicate consisting of RNA extracted from two *Daphnia* individuals. Barcoded samples from the 4 treatments within each clone were pooled together into 3 replicate libraries, purified separately, and pooled together immediately before adding the sequencing adapter. Libraries were then sequenced using Oxford Nanopore MinION for 24-48 hours per sequencing run, obtaining 2-4 Gb of reads in each run.

### RNAseq data analysis

Basecalling and reads filtering, demultiplexing, trimming, and mapping were accomplished using ONT Guppy software (ver. 3.6). *Daphnia magna* reference transcriptome 3.0 (D.Ebert and P.Fields, personal communication) was used as a reference. For the purpose of this analysis, the reference transcriptome containing only the longest isoform for each gene was used, with a total of 33,957 transcripts, of which at least one read mapped to 22,445 transcripts. Transcripts were then filtered to retain only those that contained at least 72 reads across all samples, resulting in a set of 6050 transcripts retained for further analysis.

As each library and each sequencing run consisted of 3 biological replicates of each of 4 combinations of CMIH and ASH treatments from a single clone, clones were fully confounded with replicate library preparation and sequencing runs. The advantage of this design is that each library preparation contains a balanced set of all 4 treatments with a clone-library replicate combination that can either be used as a random block effect in a 3-way ANOVa, or pulled with biological replicates for Likelihood Ratio tests (see below). The disadvantage of this design is the lack of the ability to test for the difference among clones, untangling the variance among clones from random variance among library preparation and sequencing runs.

Differential expression was analyzed using Likelihood Ratio tests (LRT) in DESeq2 (Love et al. 2014) and by 3-way ANOVa with RPKM as the response variable and CMIH treatment, ASH treatment, and habitat of origin as fixed effects and clones as a random block effect, implemented in JMP (ver. 16, SAS Institute 2021). In DESeq2 analysis, log fold change noise was shrunk by the *apeglm* algorithm for the Wald test (Zhu et al. 2019). DESeq2 analysis was conducted separately for full data, for the two habitats of origin separately, and for the analysis of the CMIH factor for the subset of samples not exposed to ASH treatment (ASH=Control). For LRT analysis, the reduced model consisted of all factors except the factor being tested and all its interactions. Wald tests and LRTs yielded similar results; with only LRT results reported. An arbitrary adjusted p-value of padj<0.1 was chosen as the cut-off for reporting differential gene expression in any given transcript. For 3-way ANOVAs, a false discovery rate (FDR) procedure was implemented for the multiple test correction, again with the cut-off of FDR<0.1. Details of statistical analyses are summarized in Supplementary Table 2, and both R and JMP scripts are available in the Supplementary materials.

Enrichment analysis was conducted separately for lists of transcripts with a possible significant effect for each of the main effects (CMYH, ASH, habitat type) and interactions. Two sources of gene lists and two types of annotation data were used. Gene lists were obtained by selecting transcripts with an uncorrected p<0.01 separately in the DESeq2 LRT analysis or in 3-way ANOVAs. The results of these analyses were similar, and only the LRT enrichment results will be reported. The two approaches to the annotation data were: a closed, non-overlapping list of transcripts with functions a priori known to be hypoxia related; and an open, overlapping list of GO categories. Firstly, we constructed a non-overlapping list of annotation terms that characterized pathways and functions a priori known to play a role in hypoxia response. The only deviations from the non-overlapping principle were the fused genes containing vitellogenin and superoxide dismutase domains (Kato et al. 2004), which appeared both in the “vitellogenins” and “antioxidant pathway” gene lists. The source of annotation was a combination of descriptions obtained from the *D. magna* genome annotation available at (http://arthropods.eugenes.org/genepage/daphniamagna/Dapma7bEVm000001t1; Gilbert 2002) and annotations obtained for the *D. magna* 3.0 transcriptome by blast2GO (Götz et al. 2008) and PANTHER (Mi et al. 2013) software. Secondly, we used all GO annotations generated by blast2GO and PANTHER to construct an open list of all GOs; this list was filtered to include only GOs represented in the reference transcriptome by at least 10 transcripts. In both cases, the Fisher exact test was used to test for enrichment, testing the hypothesis that a given pathway, function, or GO had a higher than random representation in the list of candidate genes (left-handed FET).

## Results

### Lifespan in chronic mild intermittent hypoxia

CMIH treatment provided a life-extending effect in clones originating from intermittent habitats but not in clones originating from permanent habitats (Fig. 1; Table 1). The median lifespan increased from 65 days (60-73, 95% CI) to 97 days (85-106, 95% CI) in clones from intermittent habitats. In the clones from permanent habitats, the median lifespan decreased slightly and not statistically significantly from 82 days (76-89 95%CI) to 70 days (64-77 95%CI). The proportional hazards test of the habitat x hypoxia interaction term had a P<0.0028; while the clone x hypoxia interaction was not significant (Table 1).

**Fig. 1.**
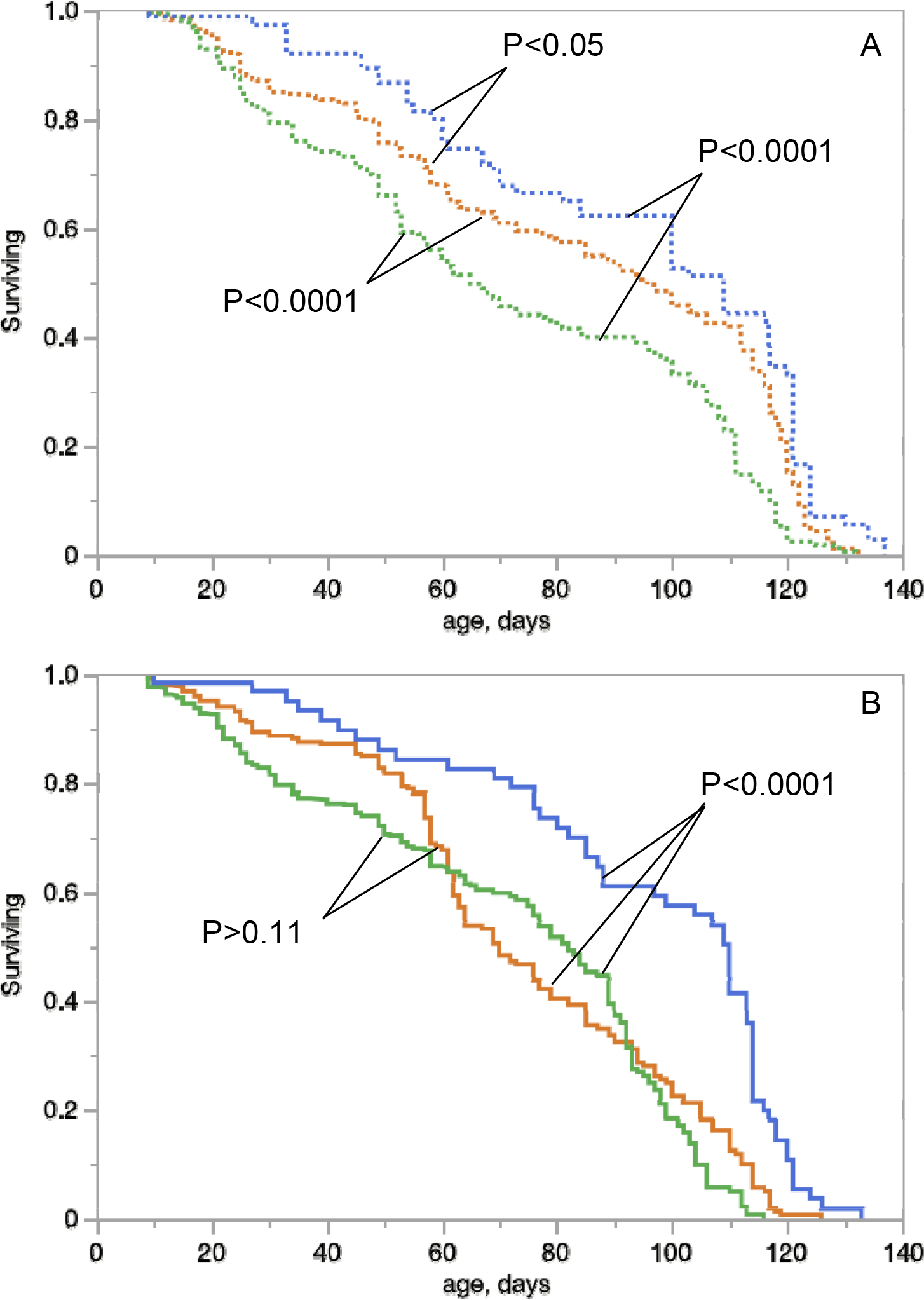
Lifespan of *Daphnia* from intermittent habitats (A, dashed lines) and permanent habitats (B, solid lines) in normoxic conditions (8 mgO_2_/L, green), chronic mild intermittent hypoxia (4 mg O_2_ /L twice daily; orange), or after switch from 4 to 8 mgO_2_/L at day 30 (blue). P values for Log-rank test for survival differences between groups are shown. See Table 1 for detailed survival analysis. See Supplementary Fig. S1 for the same data grouped by hypoxia conditions rather than by habitats of origin.

The switch from CMIH to normoxic treatment provided further extension of lifespan in all clones (Table 1), increasing median lifespan to 109 days (100-117 95%CI) in clones from intermittent habitats and to 110 days (88-113 95%CI) in clones from permanent ones. Switch vs. constant treatments contrasts revealed that the effect of the switch was significant in clones from intermittent habitats in the switch vs. normoxia contrast (model parameter = 0.384 ± 0.0694) but not in the switch vs. CMIH treatment contrast (model parameter = -0.0214 ± 0.066). In clones from permanent habitats, both contrasts were significant (0.261±0.071 and 0.359 ± 0.074 in switch to CMIH and switch to control contrasts, respectively).

### Body size, fecundity, feeding rate, and respiration rate in CMIH vs. control

Chronic mild intermittent hypoxia had a small effect on *Daphnia* body size at maturity, but there was a significant interaction effect (Fig.2A, Supplementary Table S3): clones from permanent habitats (which had lower size at maturity in controls) became even smaller in hypoxic conditions, while those from intermittent habitats became larger. *Daphnia* reared in CMIH had a lower feeding (filtering) rate than their normoxic control counterparts (Fig.2B, Supplementary Table S3). This result held regarding whether the feeding rate was normalized by the wet weight of *Daphnia* or not. In particular, CMIH- reared *Daphnia* from the GB clone had an almost undetectably low feeding rate in this experimental setup. In contrast, respiration rate did not differ between CMIH and control *Daphnia* (Fig.2C, D, Supplementary Table S3), either with an oxygen assay conducted with an initial oxygen concentration of 8 mg/L (Fig.2C), or with *Daphnia* from each CMIH treatment tested with the initial oxygen concentration matching their rearing conditions (Fig.2D).

**Fig. 2.**
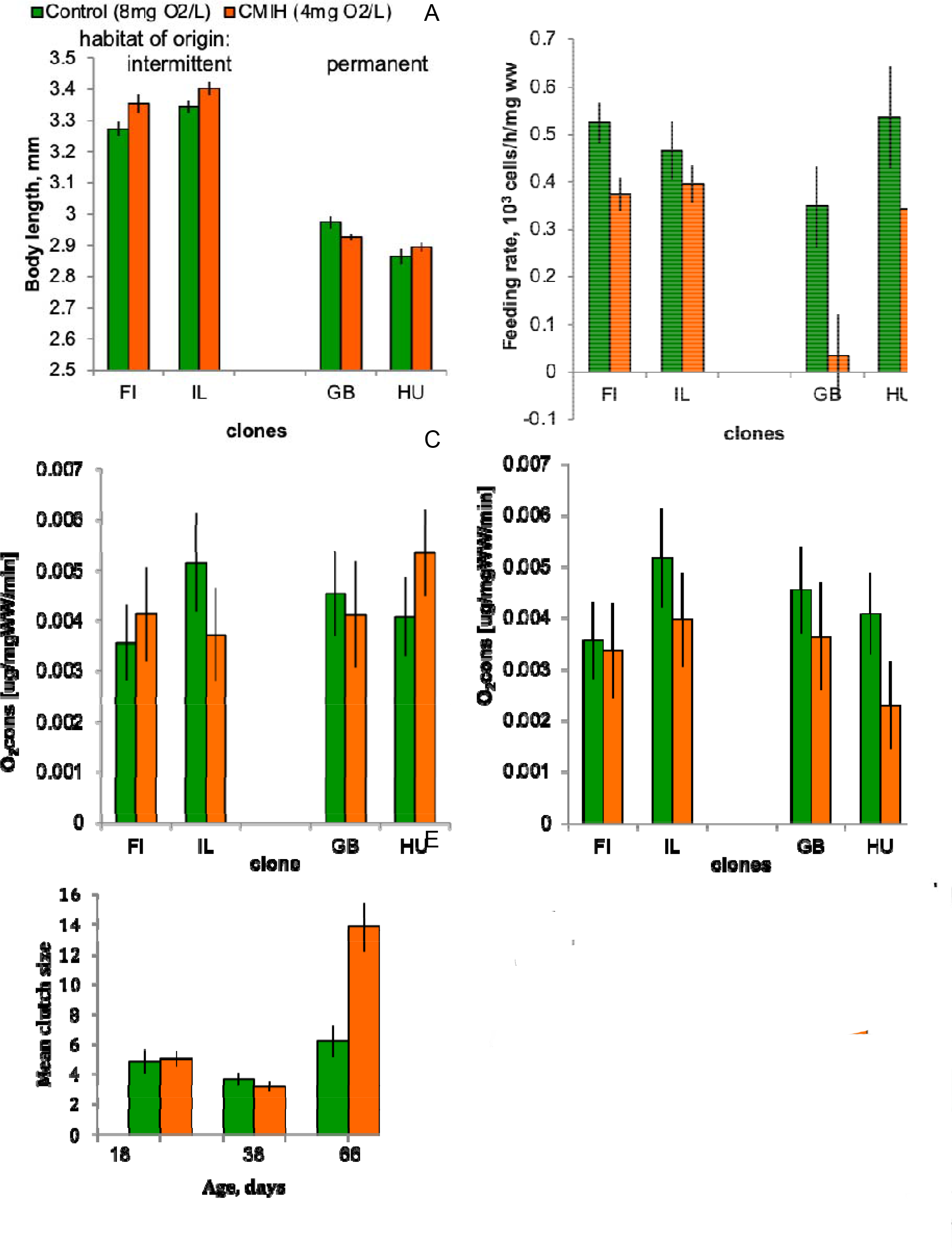
Body length at maturity (A), feeding ra (B), respiration rate (C, D) and fecundity (E) i four *Daphnia* from intermittent (FI, IL) and permanent (GB, HU) habitats reared either in normoxic conditions (8 mgO_2_/L, green) or chronic mild intermittent hypoxia (4 mg O_2_ /L twice daily; orange). Respiration rate is measu either at an initial O_2_ concentration of 8 mg/L or at an initial O_2_ concentration equal to that in the rearing conditions (D). The bars here and below are standard errors. See Supplementary Table S3 for statistics.

Finally, early fecundity did not differ between CMIH and control treatment, but at the age of 66 days, CMIH-reared *Daphnia* showed significantly higher clutch sizes than those in the control (Fig. 2E, Supplementary Table S3).

### Survival in severe acute hypoxia

Contrary to the predictions of protective or hormesis-like acclimatory effects of CMIH on hypoxia tolerance, *Daphnia* from the CMIH treatment showed lower acute hypoxia tolerance in ASH trials than their normoxia-reared counterparts (Fig. 3, Table 2). Clones from intermittent habitats had a significantly higher ASH tolerance than those from permanent ones (Fig. 3, Table 2), with clones within habitat types being only marginally significantly different from each other.

**Fig. 3.**
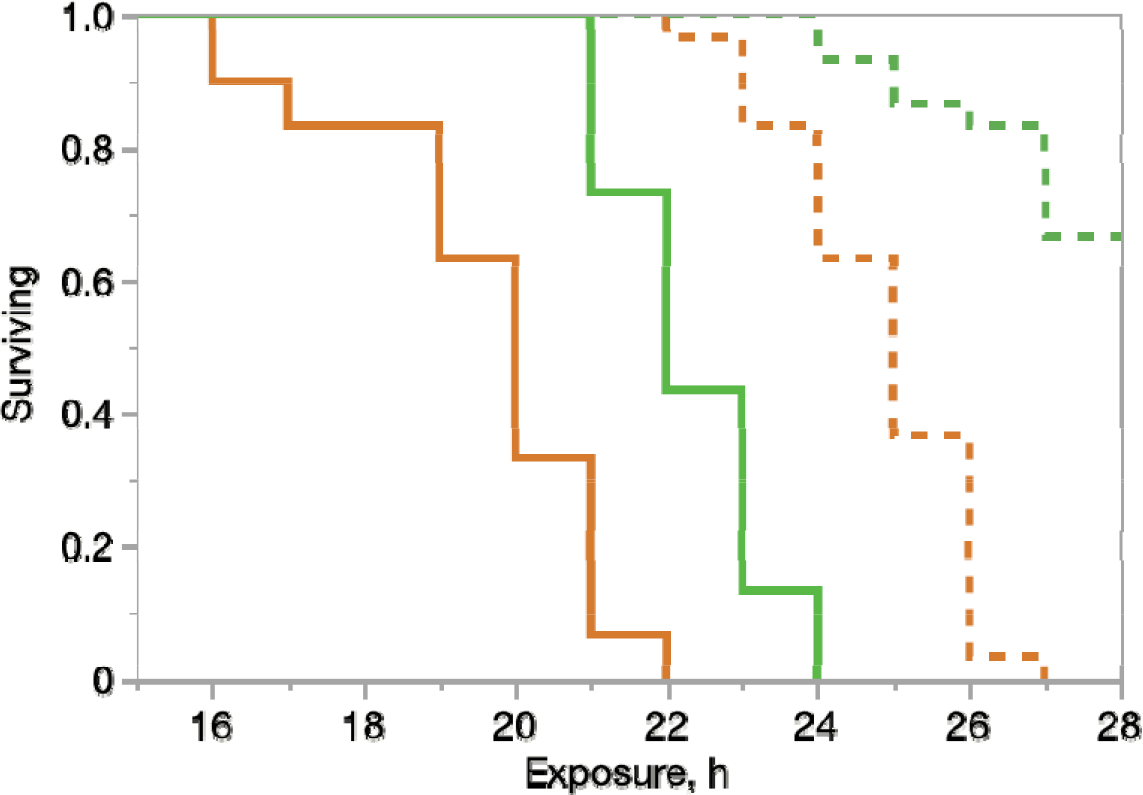
Survival of *Daphnia* from intermittent habitats (dotted lines) and permanent habitats () exposed to normoxia (green) or chronic mild intermittent hypoxia (4 mg O_2_ /L twice orange). See Table 3 for detailed survival analysis.

**Table 2.** Likelihood Ratio proportional hazards tests of the effects of habitat type, clones nested within habitats, chronic hypoxia, and their interactions on Daphnia acute hypoxia survival (Fig.4).

| Source | DF | Log Ratio $\chi^2$ | P value |
| --- | --- | --- | --- |
| habitat | 1 | 149.5 | <0.0001 |
| clone[habitat] | 2 | 5.69 | 0.058 |
| CMIH | 1 | 66.7 | <0.0001 |
| CMIH *habitat | 1 | 0.199 | 0.6553 |
| CMIH *clone[habitat] | 2 | 5.39 | 0.0674 |

### Lac/Pyr ratio

As expected, *Daphnia* reared in CMIH conditions showed an elevated Lac/Pyr ratio relative to normoxic controls (Fig. 4, Table 3).This difference was especially noticeable in older (55–60-day old) *Daphnia* (Fig. 4A), and was primarily due to a decrease in pyruvate concentration (Fig. 4C; Supplementary Table S4) rather than an increase in lactate concentration (Fig. 4B; Supplementary Table S4). Contrary to the prediction of decreased NAD+ availability with age (i.e., an increase in lactate concentration with age), the Lac/Pyr ratio dropped significantly between age classes of 15-20 and 55-60 days. This drop, again, was largely explained by the increased pyruvate concentration (Fig. 4C; Supplementary Table S4). *Daphnia* reared in the CMIH condition showed no change in Lac/Pyr ratio with age, with a slight increase in both protein content-normalized lactate and pyruvate concentrations with age, resulting in a significant (P<0.0001) CMIHxAge interaction (Table 3).

**Table 3.** Three-way ANOVA of the effects of CMIH, age, and clones on the Lac/Pyr ratio (Fig. 3 and S2). “Plate” is a random block effect.

| Source | DF | SS | F Ratio | Prob > F |
| --- | --- | --- | --- | --- |
| CMIH | 1 | 252.8 | 51.31 | <0.0001 |
| clone | 1 | 21.6 | 4.38 | 0.0389 |
| CMIH*clone | 1 | 1.26 | 0.26 | 0.61 |
| age | 1 | 137.4 | 27.89 | <0.0001 |
| CMIH *age | 1 | 112 | 22.75 | <0.0001 |
| clone*age | 1 | 0.21 | 0.04 | 0.84 |
| CMIH *clone*age | 1 | 0.06 | 0.01 | 0.92 |
| plate | 1 | 107.8 | 21.88 | <0.0001 |
| Error | 103 | 507.79 |  |  |

**Fig. 4.**
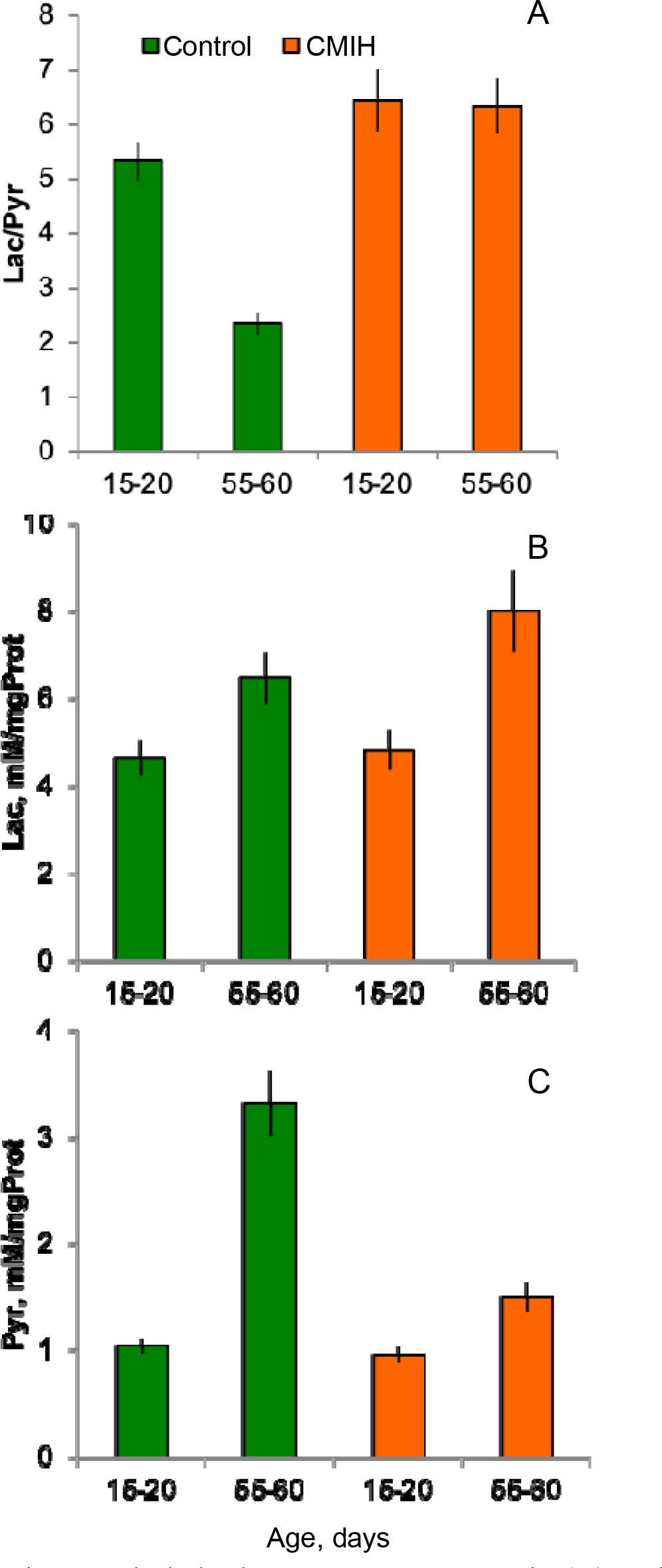
Whole body lactate/pyruvate ratio (A) and protein-normalized lactate and pyruvate concentrations (B, C) in young (15-20 days) and moderately aged (55-60 days) *Daphnia* reared at either normoxic control (green) or CMIH (orange) conditions. See Table 3 and Supplementary Table S4 for statistics, and Supplementary Fig. 2 for data separately for each clone.

**Table 4.** Differences in mitochondrial membrane potential (ΔΨ_m_) among tissues, between CMIH treatments (CMIH vs. control), between habitat of origin (intermittent vs. permanent), and between clones nested within habitat type (Fig. 5). Tukey test for differences among clones (P<0.05): all pairwise comparisons of clones across habitat types significant, except IL vs. HU (P>0.25).

| Source | DF | SS | F Ratio | Prob > F |
| --- | --- | --- | --- | --- |
| Tissue | 3 | 73617.2 | 195.5 | <0.0001 |
| CMIH | 1 | 1129.7 | 9.00 | 0.0035 |
| Habitat | 1 | 3773.9 | 30.06 | <0.0001 |
| CMIH *Habitat | 1 | 670.3 | 5.34 | 0.0232 |
| CMIH *Tissue | 3 | 294.0 | 0.78 | 0.508 |
| Habitat*Tissue | 3 | 590.6 | 1.57 | 0.203 |
| CMIH *Habitat*Tissue | 3 | 29.0 | 0.077 | 0.9722 |
| Clone[Habitat] | 2 | 1142.3 | 4.55 | 0.0132 |
| Clone*Tissue[Habitat] | 6 | 462.1 | 0.61 | 0.7189 |
| Clone* CMIH[Habitat] | 2 | 1659.6 | 6.61 | 0.0021 |
| Error | 86 | 10796.1 |  |  |

### Mitochondrial membrane potential

ΔΨ_m_ was slightly higher in clones from intermittent habitats than in clones from permanent habitats across all four studied tissues (Fig. 5, Table 5). Furthermore, clones from intermittent habitats tended to reduce their ΔΨ_m_ in the CMIH treatment more strongly than clones from permanent habitats, although individual pairwise comparisons were not significant in post-hoc Tukey test. As a result, there was a significant difference between clones from the two habitat types in control, but not in CMIH treatment, at least in the epipodite and the brain. There was therefore no evidence that hypoxia-tolerant genotypes showed any ΔΨ_m_ patterns specifically in CMIH conditions that would be different from those of hypoxia-sensitive ones or correlate with the extended lifespan.

**Fig. 5.**
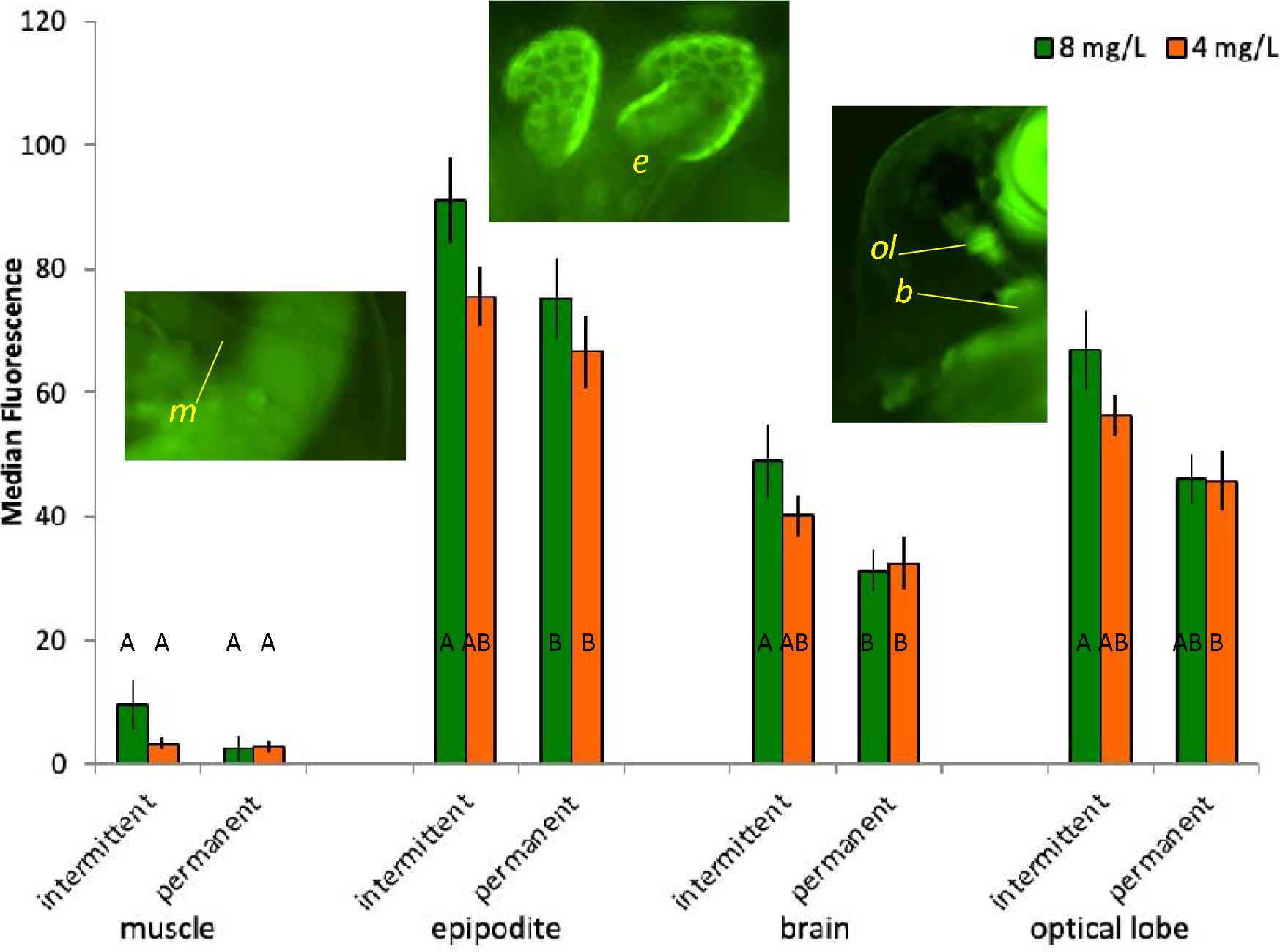
Median rhodamine-123 fluorescent used as a measure of mitochondrial membrane potential in antenna-driving muscle (m), epipodite (osmoregulation/gas exchange organ, e), brain (b), and optical lobe (ol) in *Daphnia* from either intermittent or permanent habitats reared either in normoxic (8 mg O_2_/L, green) or CMIH (4 mg O_2_/L twice daily, orange) conditions. Letters on the bars are the results of Tukey test conducted for each tissue separately. See Table 4 for statistical analysis.

**Table 5.**
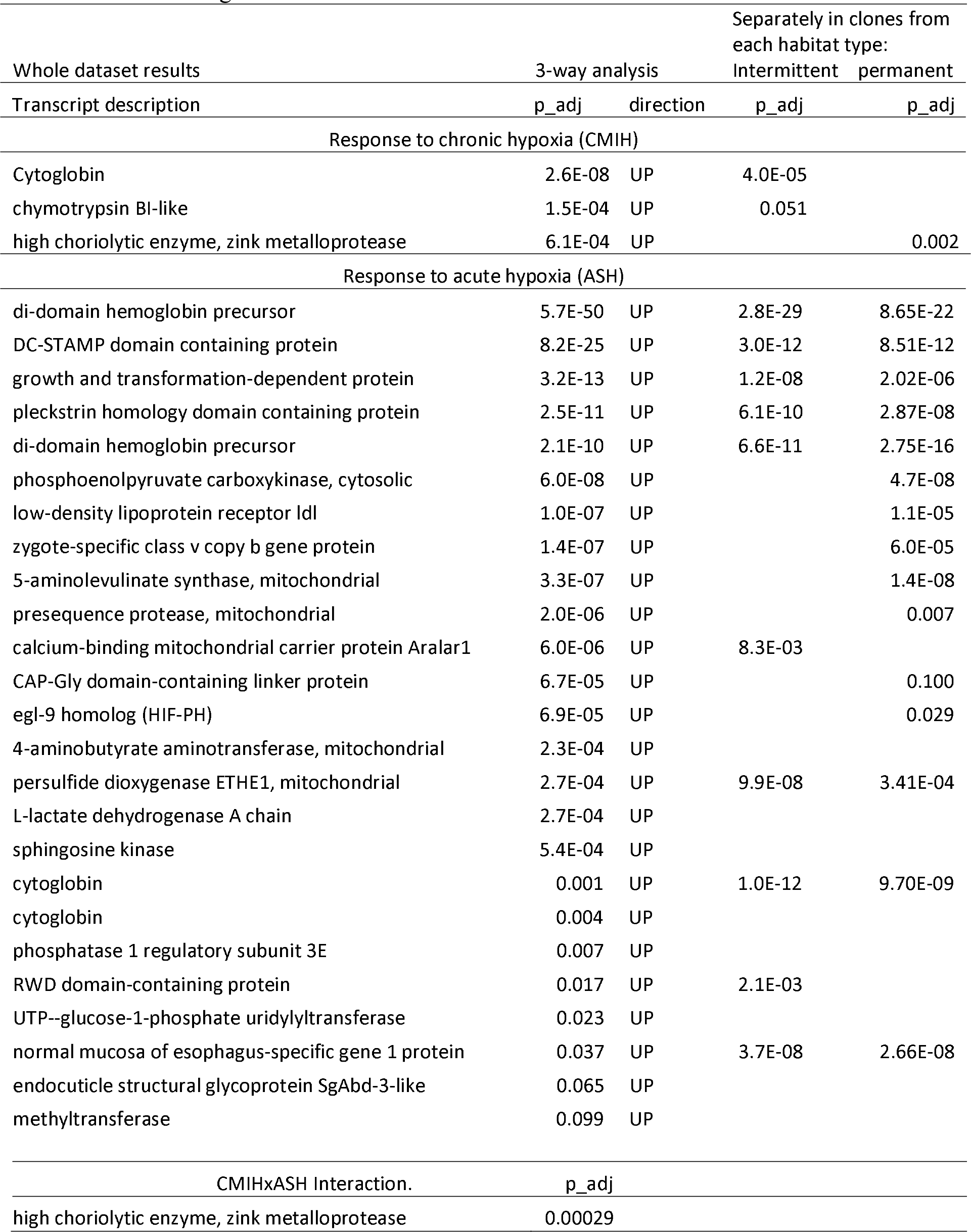
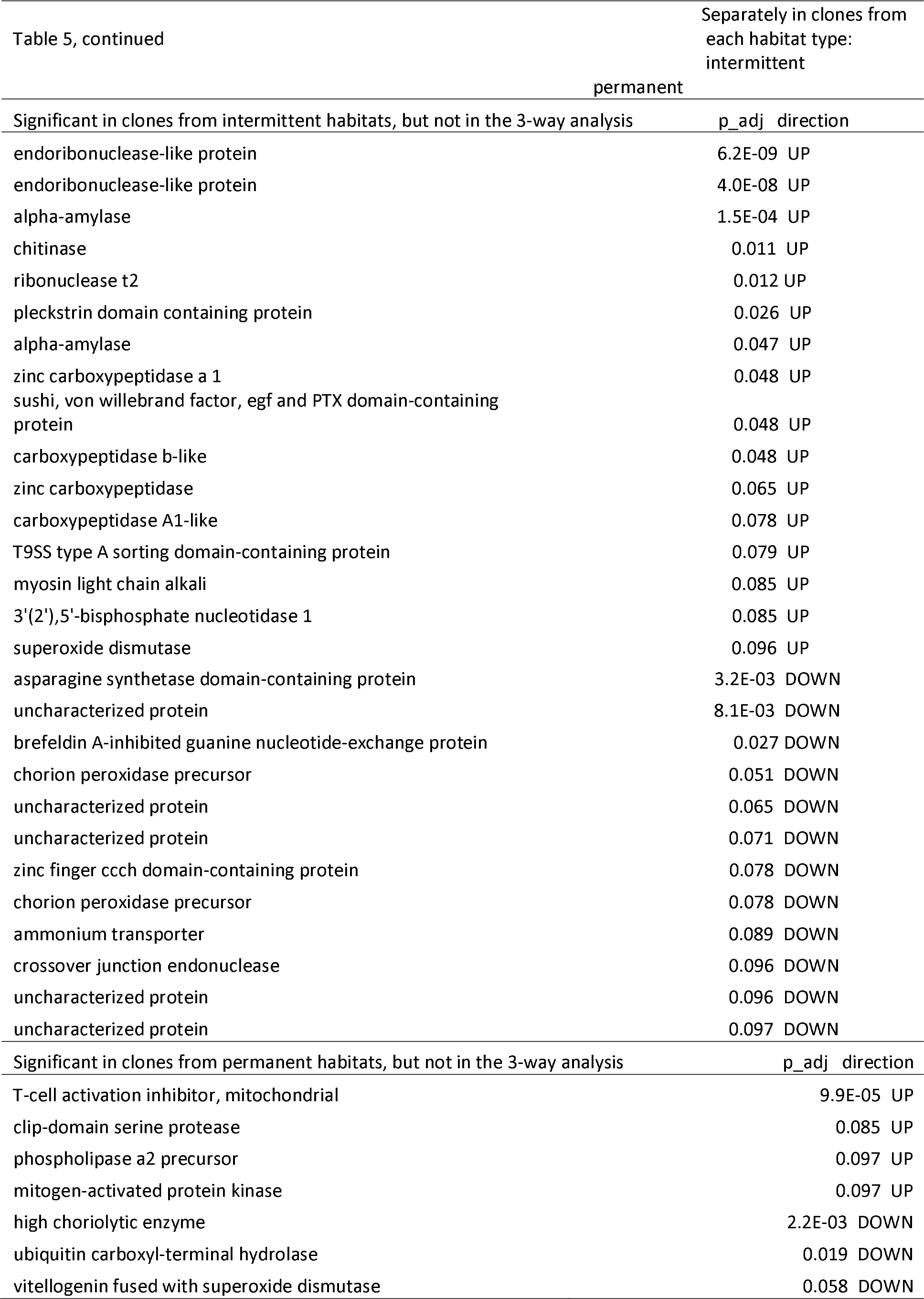
Transcripts with a significant differential expression in response to either acute or chronic intermittent hypoxia (and their interaction), in all 4 clones and in clones from intermittent and permanent habitats separately. Direction column indicate up- or down-regulation in the hypoxic treatment. Padj<0.1 used as the cut-off. Padj values not shown are >0.1, or in rare cases, the LRT <u>model failed to converge.</u>

### Differential gene expression in response to CMIH and ASH

Only three transcripts individually survived multiple test correction in the 3-way analysis of responses to CMIH, both in the LRT analysis (Table 5) and in 3-way ANOVAs with clones as the random block effect, all three showing up-regulation in the CMIH treatment. This result did not change when the CMIH effect was analyzed within the ASH=Control subset (i.e., in *Daphnia* never exposed to acute hypoxia only) or in the analyses with each habitat of origin type separately (data not shown). These transcripts were (Table 5, Supplementary Fig. S4 C,J): one of the cytoglobin paralogs (a homolog of mammalian CYGB globin known to protect against hypoxia, probably by facilitating diffusion of oxygen through tissues and scavenging nitric oxide or reactive oxygen species; Trent & Hargrove 2002) and two proteases, a chymotrypsin BI-like protease and a high choriolytic enzyme zinc metalloprotease (with no known hypoxia-related functions). Additionally, the latter transcript was the only one showing a significant CMIHxASH interaction (Table 5, Supplementary Fig. S4).

A significantly longer, fully non-overlapping list of transcripts showed a significant response to ASH treatment, in all versions of the analysis, 3-way LRT analysis, LRT analysis separately in clones from the two habitat types (Table 5; Supplementary Fig. S4) and in the 3-way ANOVAs, again, all showing up-regulation in ASH condition. Prominent in this list were the di-domain hemoglobin precursors well characterized for their plastic response to hypoxia (Zeis et al. 2009; Zeis 2020), as well as cytoglobins and mitochondrial 5-aminolevulinate synthase critical for heme synthesis (Supplementary Fig. S4 A, B, E), and, not surprisingly, one of the homologs of egl-9 (HIF prolyl hydroxylase) present in *Daphnia* (Supplementary Fig. S4 F). Other transcripts up-regulated in ASH include those coding for rate-limiting enzymes in anaerobic metabolism such as lactate dehydrogenase (Ldh) and cytosolic phosphoenolpyruvate carboxykinase (PEPCK-C; Supplementary Fig. S4 G, H). There were also numerous ASH-upregulated transcripts with mitochondrial membrane localization and functionality, as well as many with unclear relation to hypoxia and respiration, including several proteases.

In the analysis conducted separately for clones from each of the two habitat types, many of the same transcripts showed a significant differential expression in response to the ASH treatment (Table 5) in both types of clones, including hemoglobins and cytoglobins (Table 5, Supplementary Fig. S4, A-D). Yet, many transcripts showed significant upregulation in only one of the two types of clones. In particular, 5- aminolevulinate synthase, PEPCK-C and egl-9 were significantly overexpressed in hypoxia-sensitive permanent habitat clones, but not significantly so in the hypoxia-tolerant intermittent habitat clones (Table 5, Supplementary Fig. S4, E-G). Additionally, the permanent habitat clones showed stronger up-regulation of a low-density lipoprotein receptor, zygote-specific class v copy b protein, and a mitochondrial presequence protease. Furthermore, there were several transcripts up- or down-regulated exclusively in this type of clones, but not in the whole dataset analysis, including, notably, the one coding for the mitogen-activated protein kinase (MAPK) homolog, the key regulator of a general stress signaling pathway.

A different set of transcripts changed expression in the ASH treatment in the hypoxia-tolerant, intermittent habitat clones (Table 5). Among them there were: calcium-binding mitochondrial carrier protein Aralar1, several endoribonucleases, several – α amylases, a chitinase, numerous carboxypeptidases, and one of several paralogs of SOD.

### Functional enrichment of differentially expressed genes: a priori defined gene set with hypoxia-related functions

Targeted analysis of such transcripts revealed a significant enrichment of several a priori defined pathways or functions among transcripts with uncorrected p-values in one or several types of LRT analysis (Table 6). Antioxidant pathways (represented mostly by SOD paralogs) as well as globins showed enrichment in LRT analysis of all factors in the model. Respiratory, ECH, and ATP synthesis pathways did so in the analysis of the ASH effect and, with a less significant FDR value in the CMIHxASH interaction analysis.

**Table 6.**
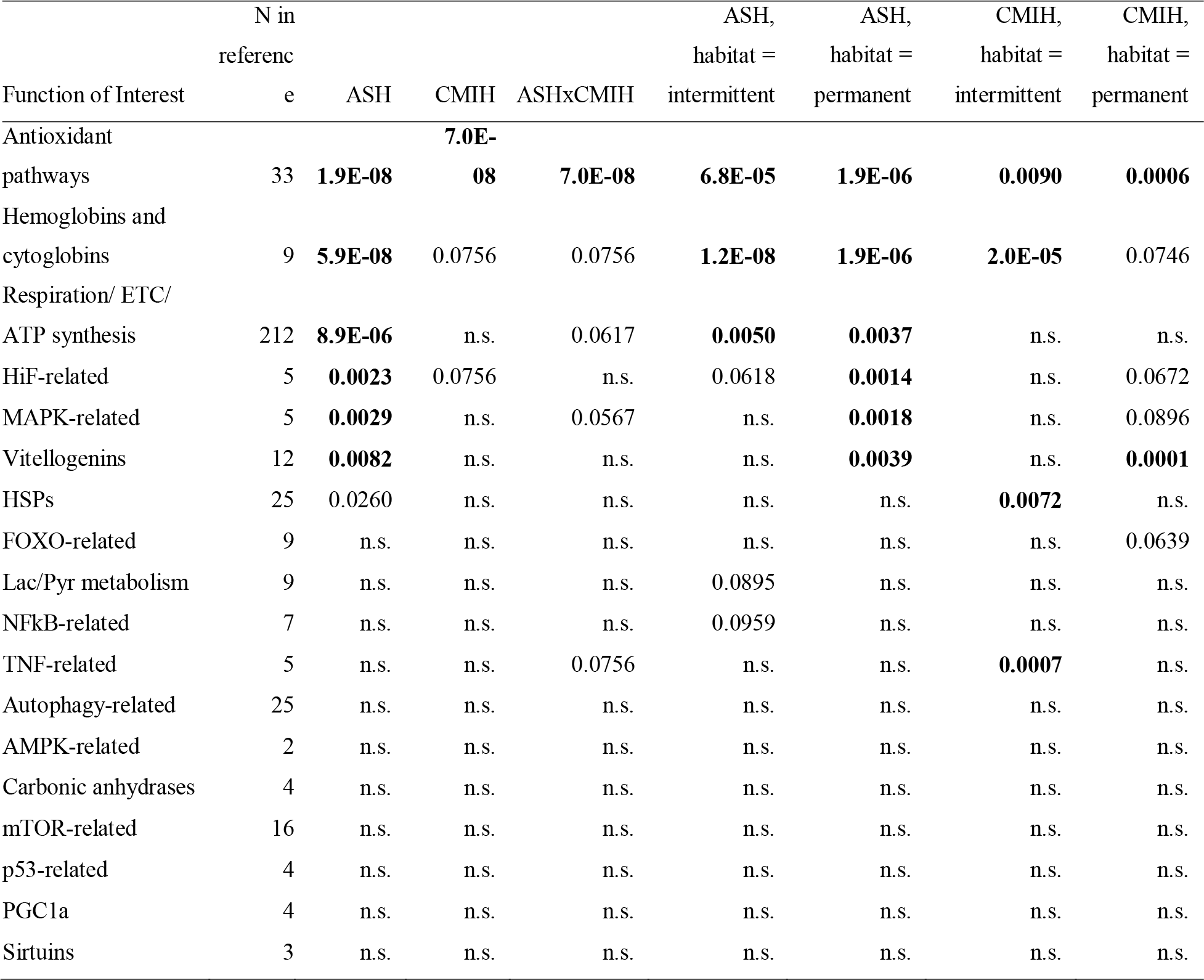
Representation of a priori hypoxia-related functions of interest in gene lists with uncorrected p-values <0.01 for CMIH, ASH and their interactions in the whole data and estimated for clones originated separately from intermittent and permanent habitats. Values are FDR-adjusted Fisher exact test p-values of significant over-representation (bold: FDR<0.01).

Other targeted functionalities were more specific to individual analyses. General hypoxia-related (such as the egl-9 paralog), MAPK-related proteins and vitellogenins appeared to be specific to clones from permanent habitats. In contrast, HSPs, Lac/Pyr metabolism-related proteins, as well as NFkB-related and TNF-related proteins were enriched in the analyses specific to clones from intermittent habitats. It appears that clones from different habitats engaged radically different repertoire of plastic transcriptional responses to either CMIH or ASH treatments. Notably, none of the other hypoxia-related pathways showed any enrichment in any of the analyses, including FOXO-, autophagy, AMPK, mTOR, and p53-related pathways or the sirtuins (Table 6).

Similar results were obtained using gene sets with uncorrected p-values in the 3- way ANOVAs with clones/library prep blocks as a random factor (results not shown).

### Functional enrichment of differentially expressed genes: open list of GOs or pathways

Few GOs or pathways identified by blast2go and PANTHER scans showed any significant enrichment among any gene lists with possible differential expression (although many showed a significant under-representation; results not shown). There were several possible exceptions overrepresented in a single gene set: those with an uncorrected p-value <0.01 for the ASHxhabitat type interaction in the 3-way ANOVA tests. These GOs included various nested GO categories already revealed by the previous analysis, such as heat shock proteins or mitochondria-related GOs. There were, however, also two types of GO or pathway categories not revealed by the previous analyses: extracellular matrix structural proteins (PC:00103) and extracellular region (GO:0005576), which were overrepresented due to the high occurrence of cuticular proteins (Supplementary Fig. S3, A) and central nervous system development (GO:0007417), over-represented due to a group of linked paralogous neutrophins (Supplementary Fig. S3, B), encoded by genes located in scaffold region 6F30.52-30.164. Both these groups of transcripts show up-regulation in *Daphnia* from intermittent habitats and down-regulation in those from permanent habitats. None of the transcripts reached a significant FDR for the ASH x habitat interaction, but both GOs are over-represented with FDR<0.0001.

### Multidimentional analysis of differentially expressed transcripts

Principal component analysis of 48 samples (2CMIH x 2ASH x 4clones/libraries x 3 biological replicates) in the space of 587 transcripts with at least one uncorrected p<0.01 is shown in Fig. 5. Several patterns are apparent from this analysis. Firstly, there were strong differences among confounded clone/library replicates. Secondly, within each such replicate, there was a separation between ASH and control biological replicates (Fig 6, triangles vs. circles), but not between CMIH vs. normoxic control (orange vs. green), consistent with fewer transcripts reaching significant differential expression with multiple test correction. Thirdly, this separation was significantly stronger in the clones from intermittent habitats (FI and IL) than in those from permanent habitats (GB and HU): all 3 first principal components showed a significant separation of ASH vs. control in FI and IL, controlling for random differences between clones/libraries, but only the third principal component showed such separation in the GB and HU clones analyzed separately (Supplementary Table 5).

**Fig. 6.**
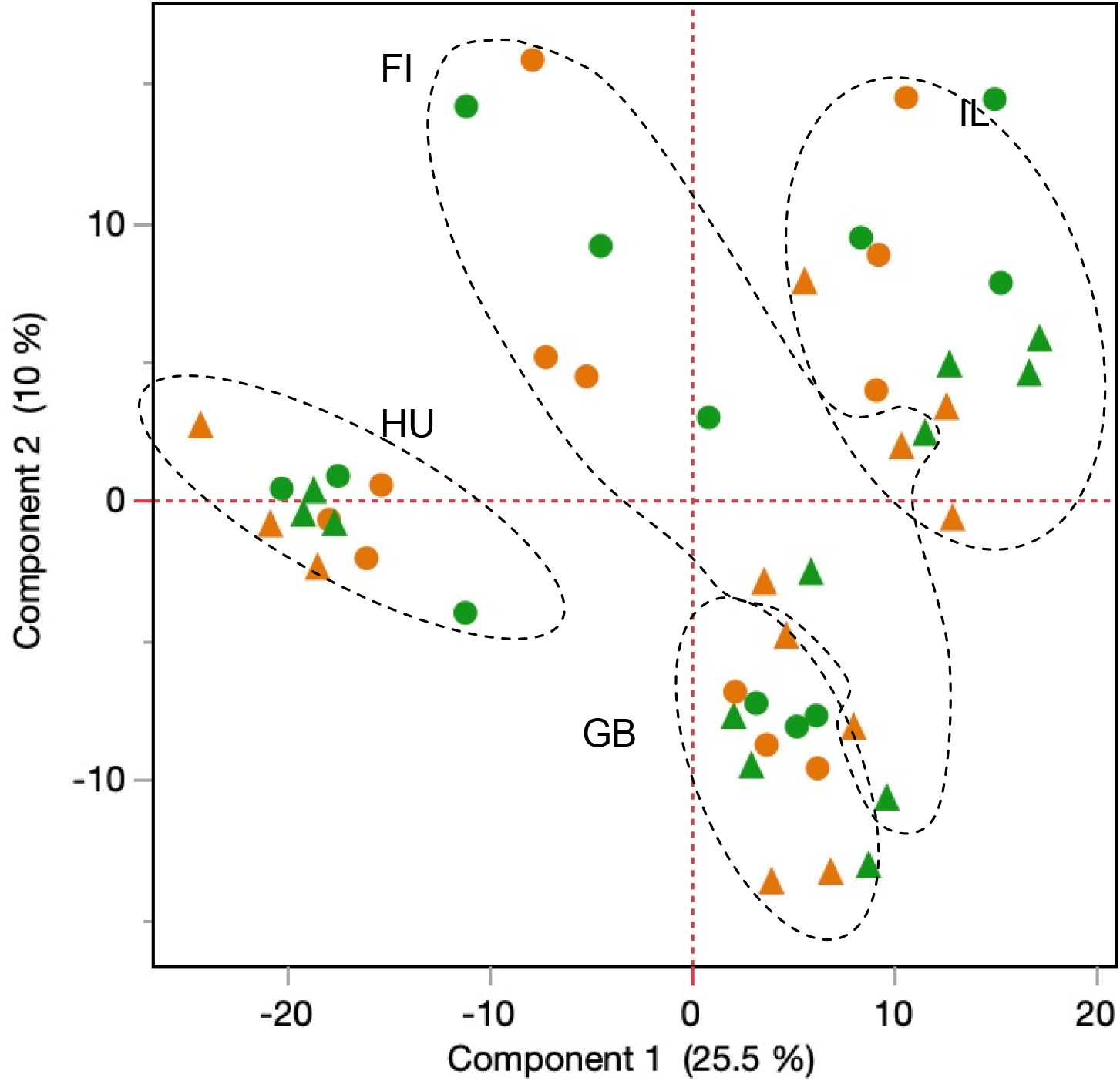
PCA of 48 samples (2CMIH x 2ASH x 4clones/libraries x 3 biological replicates) in the space of 587 transcripts with at least one uncorrected p<0.01. Colors as on previous figures, namely green: CMIH control, orange: CMIH treatment. Circles: ASH control, triangles: ASH treatment. Ellipses encircling clone/library blocks drawn by hand.

Finally, while the hypoxia-tolerant clones from intermittent habitats (FI and IL) were somewhat separated from the hypoxia-sensitive ones from permanent habitats (GB and HU), the direction of change from control to ASH treatment did coincide with this difference. Exposed to acute hypoxia, the sensitive clones did not change expression patterns to match the tolerant clones; rather, the hypoxia-tolerant clones exposed to acute hypoxia became more transcriptionally similar to the sensitive ones (Fig. 6). In other words, most of the differential expression did not reflect adaptive plasticity and/or canalization of transcriptional phenotype in the hypoxia-tolerant clones.

## Discussion

### Chronic mild intermittent hypoxia extends lifespan in Daphnia from intermittent habitats

We observed an expansion of lifespan in CMIH treatment in clones from hypoxia-prone, intermittent habitats, but not in clones from permanent habitats that rarely experience hypoxic conditions. While this can be readily interpreted as local adaptation, we did not see any manifestation of it on the transcriptional level other than the up- regulation of one of the cytoglobin paralogs in the clones from intermittent habitats, but not in those from permanent habitats. We must therefore conclude that, with the exception of the cytoglobin expression, this intriguing opposite directionality of CMIH on lifespan is not based on transcriptional profile differences among genotypes. Likewise, we did not observe any differences in respiratory phenotypes (respiration rate, lactate/pyruvate ratio, and mitochondrial membrane potential, ΔΨ_m_) interpretable as possible causes of lifespan extension in intermittent habitat clones in mild hypoxia.

Respiration rate showed no significant differences at all; lactate/pyruvate ratio and ΔΨ_m_ showed differences between CMIH treatment and control consistent with reduced aerobic respiration in CMIH, but no differences between clones with and without lifespan extension effect. Higher ΔΨ_m_ in shorter-lived clones from intermittent habitats apparent in the normoxic control is consistent, at least for the epipodite ΔΨ_m_, with the data reported in Anderson et al. 2022, as well as with data on higher respiratory metabolism in short-lived genotypes of *C.elegans* (Feng et al. 2001; Lee et al. 2010).

The most significant expansion of lifespan occurred in *Daphnia* switched from the CMIH treatment to normoxia at the age of 30 days. Because this switch treatment included only one replicate tank, this result is highly speculative. In particular, the difference between switch and CMIH treatments in clones from intermittent habitats is suspect because it became apparent (Fig. 1) even before the switch, and thus represents random variation. It was also not significant in the switch vs. CMIH contrast for intermittent habitat clones, in contrast to all other contrasts. With this caveat in mind, if we were to believe that early life mild intermittent hypoxia provides a greater life-extending effect than lifetime exposure, this may indicate that protective hypoxia-induced pathways must be activated before aging-related damages have started to accumulate or before the mid-life transcriptional profile takes shape.

The late-life increase in fecundity observed in CMIH treatment across all clones is intriguing. We have documented such late-life fecundity spikes in several independent experiments (Yampolsky, Anderson, Lowman et al., in preparation), but the difference between CMIH and normoxia control observed here requires further confirmation, as it is based on a single day egg count and therefore may be spurious.

### Chronic mild intermittent hypoxia does not increase acute hypoxia tolerance

Mild levels of stress are often supposed to acclimate or prepare organisms to tolerate higher, perhaps lethal levels of the same or related stress factor. However, the hormesis principle (Mattson 2007; Maynard 2011; Berry & López-Martínez 2020), should not be taken for granted (Axelrod et al. 2004). For example, in *Daphnia,* while adaptive effects of moderate heat acclimation have been well established (Yampolsky et al. 2014; Burton et al. 2020), the present study does not show a similar effect for moderate hypoxia. To the contrary, *Daphnia* reared in mild intermittent hypoxia showed lower survival in acute trials. Sometimes what does not kill us just makes us weaker. That is not to say that no acclimatory changes occur in hypoxia: beneficial effects of mild hypoxia in heat- and oxidative stress tolerance have been documented in *Daphnia* (Klumpen et al., 2017; Coggins et al. 2017). However, this plasticity appears to convey no benefit in terms of surviving in acute hypoxia.

### Adaptive transcriptional responses to acute hypoxia

We observed several readily interpretable changes in acute hypoxia treatment.

Apart from hemoglobin precursors and mitochondrial 5-aminolevulinate synthase, a rate-limiting enzyme in the heme synthesis pathway, transcripts that were up-regulated in the ASH condition included two key enzymes in pyruvate metabolism: Ldh and PEPCK-C. Ldh accomplishes anaerobic fermentation recycling NAD+, which allows ATP production to continue via glycolysis. PEPCK-C is the rate-limiting enzyme in early stages of gluconeogenesis and thus is critical for ATP production in gluconeogenesis- dependent tissues such as muscles via the Cori cycle. Up-regulation of UTP-glucose-1- phosphate uridylyltransferase (UGP) is also consistent with these metabolic changes, assuming that in the shortage of glucose-6-phosphate and glucose-1-phosphate, it can replenish glucose-1-phosphate from glycogen storage. Finally, up-regulation of calcium-binding mitochondrial carrier protein Aralar1 is also consistent with the apparent importance of gluconeogenesis, as its function is the transport of aspartate from the mitochondria to the cytosol, where it can be converted to oxaloacetate, the starting substrate for gluconeogenesis.

Up-regulation of gluconeogenesis by hypoxia is paradoxal. The connection between hepatic hypoxia and gluconeogenesis has been recently documented in mammals (Ramakrishnan et al. 2016; Ramakrishnan & Shah 2017) and fish (Sun et al. 2020), but the direction of change is exactly the opposite: hypoxia down-regulates gluconeogenesis through three different pathways, all critically linked to HiF2α Yet, in a recent study in *C.elegans,* up-regulation of PEPCK-C in hypoxia has also been observed (Vora et al. 2021), indicating that up-regulation of gluconeogenesis in hypoxia may be a widespread phenomenon in invertebrates. What may be the functional significance of this reverse effect of hypoxia on gluconeogenesis? We believe that the key difference is in the heterogeneity of oxidation levels among tissue that exists in vertebrates. Notably, the liver is in a normoxic state during fasting and becomes hypoxic after refeeding when central blood flow is redirected towards the gastrointestinal tract (Ramakrishnan et al. 2016). This signals down-regulation of the glucagon pathway and thus gluconeogenesis in the liver, contributing to the maintenance of blood glucose homeostasis. In animals like *C.elegans* or *Daphnia,* this mechanism is not likely possible due to little or no lymph circulation regulation and a small body size which probably results in a uniform oxygen availability throughout the body. In *Daphnia,* oxygen concentration in tissues shows an anterior-posterior gradient (less pronounced when hemoglobins are present in hemolymph) that reflects the general direction of circulation (Pirow et al. 2004). Thus, rather than switching between up- and down-regulation of gluconeogenesis in response to feeding/fasting phases, hypoxia up-regulates gluconeogenesis to supply glucose for glycolysis in critically important tissues such as heart and striated muscles, to provide substrates for the pentose phosphate pathway, and to replenish NADPH that can be utilized in glutathione biosynthesis (Vora et al. 2021), thus contributing to the reduction of oxidative stress.

It may be noted that transcripts up-regulated specifically in clones from intermittent habitats (low number of clones per habitat type caveat notwithstanding) are dominated by catabolism enzymes encoding transcripts: ribonucleases, amylases, and peptidases, notably carboxypeptidases, with likely catabolic functions (Schwerin et al. 2009), indicating possible adaptive significance of activating ATP-producing pathways alternative to aerobic respiration.

### Transcriptional changes in response to hypoxia are not necessarily adaptive

There were numerous transcripts responding to acute hypoxia for which there was no obvious adaptive explanation, or which, counter-intuitively, showed a stronger response in hypoxia-sensitive than in hypoxia-tolerant clones. There may be several reasons for this observation. Firstly, some of these changes may be constrained, passive downstream responses to hypoxia (cf. Yampolsky et al. 2014b) rather than adaptive ones. The nature of such responses may have to do with common regulatory pathways that are shared for entirely different reasons rather than adaptation to hypoxia, as seen, for example, in salt tolerance responses in yeast (MacGilvray 2018). Alternatively, a transcriptional or phenotypic response to stress may simply occur because stress disrupts regulatory pathways in a constrained manner, as it is known, for example, for mitochondrial quality control pathways disrupted by oxidative stress (Ambekar et al. 2021). While these non-adaptive responses can potentially be used as markers of hypoxic stress, they do not ameliorate damage caused by hypoxia. This is possibly the scenario we observe in the cases of some up-regulated proteins with no obvious hypoxia- or respiration-related functionality. The PCA analysis indicating that hypoxia-tolerant clones become more like hypoxia-sensitive ones when exposed to the ASH treatment and not *vice versa*, is suggestive that many transcriptional changes are indeed non-adaptive.

Secondly, some of the transcriptional or phenotypic changes may be occurring in the adaptive direction but are not sufficient in magnitude relative to the strength of the stress factor. This is perhaps what we observe for the transcriptional responses to CMIH of globins or antioxidant enzymes encoding genes. Despite an arguably adaptive up- regulation of these transcripts in the CMIH treatment, they, however, do not convey a higher tolerance to the ASH treatment (Supplementary Figure S4, A-D).

Finally, one may observe a phenotypic or transcriptional difference that is higher in sensitive individuals than in tolerant ones (Supplementary Figure S4). This may reflect the higher internal damage experienced by the sensitive organisms rather than the higher adaptive value of the plastic response. We hypothesize that the stronger up-regulation of egl-9 homolog (HIF-PH) by the ASH treatment in clones from permanent habitats than in clones from intermittent habitats reflects a greater degree of oxygen shortage in tissues (perhaps due to a lack of some other protective or compensatory mechanisms). Likewise, the up-regulation of MAPK, the key protein in a general stress signaling pathway, that we observe in clones from permanent, but not intermittent habitats, may be explained by the same conjecture. A similar up-regulation of MAPK has recently been observed in a relatively hypoxia-sensitive carp species (Zhou et al. 2020). We have previously demonstrated similar counter-intuitive expression patterns in *Drosophila* in response to alcohol (Yampolsky et al. 2012) and in *Daphnia* in response to heat (Yampolsky et al. 2014b).

### Functional enrichment

Enrichment analyses based on a priori expected hypoxia-related pathways and functions and on gene ontologies (Supplementary Fig. S3) produced different results. Among a priori known hypoxia-related pathways (Table 6), predictably, genes involved in antioxidant pathways, hemoglobins and cytoglobins, and HIF pathway-related genes were enriched among transcripts responding to all treatments, while membrane phosphorylation-related genes were enriched only among those responding to the ASH treatment. Other a priori known functionalities were enriched only in a subset of gene lists: for example, vitellogenins were only enriched among transcripts responding to ASH in clones from permanent habitats; HSPs and TNF pathway-related genes – only among transcripts responding to CMIH in clones from intermittent habitats (Table 6). Despite their high occurrence in the reference, the autophagy-related and mTOR-related genes showed no enrichment in any of the analyses.

None of these a priori identified functional groups were identified in the GO enrichment analysis. In contrast, two other groups of genes appeared to be non-randomly associated with hypoxia response: cuticular proteins and neurotrophins. In both these gene groups, the same pattern of ASH-by-habitat of origin interaction was observed: they were up-regulated in the ASH treatment in the clones from intermittent habitats and down-regulated in the clones from permanent ones (Supplementary Fig. S3). For cuticular proteins, one may hypothesize that hypoxia-adapted *Daphnia* from the intermittent habitats respond to hypoxia by accelerating the molting cycle – a possible mechanism to increase the uptake of oxygen in hypoxic conditions due to the higher oxygen penetrability of soft post-molt exoskeleton (Mangum et al. 1985; Peruzza et al. 2018). Peruzza et al. (2018) also observed changes in cuticular protein expression in response to hypoxia. On the other hand, it is less obvious why hypoxia-sensitive clones down-regulate cuticular proteins; perhaps it is a part of a general trend to conserve resources by down-regulating non-essential gene expression.

### Much less can be concluded from the same patterns observed in neurotrophins

First of all, this enrichment may be spurious as these genes are closely linked and may be under a common transcriptional regulation, thus representing a single independent transcriptional response, rather than six. However, if indeed not caused by chance alone, the radically different responses to acute hypoxia observed in hypoxia-tolerant and hypoxia-sensitive clones suggest an intriguing possibility that survival in hypoxia might depend on preventing neuronal death. Neurotrophins have known functions in maintaining neuron survival (Hempstead 2006) and recent data indicate that their well-characterized transmembrane p75 receptor plays a crucial role in neuronal death and survival during oxidative stress and hypoxia (Sankorrakul et al. 2021). A detailed characterization of the hypoxia response of *Daphnia* neurotrophin and neurotrophin receptor genes is needed to investigate this potentially interesting tolerance mechanism.

### Conclusions

Chronic mild intermittent hypoxia (CMIH) extended lifespan in hypoxia-tolerant, high mitochondrial membrane potential clones from intermittent habitats that are characterized by a short lifespan in normoxic conditions. It slightly shortened the lifespan of permanent habitat clones with longer normoxic lifespan. The CMIH treatment did not acclimate *Daphnia* to better survive acute severe hypoxia (ASH). Transcriptional changes associated with higher ASH survival include genes in pathways related to oxygen transport and storage, lactate and pyruvate metabolism, gluconeogenesis, and, specifically in hypoxia-tolerant clones, catabolism pathways, cuticulum proteins, and neurotrophins, indicating roles of anaerobic respiration, gluconeogenesis, modulating molting cycle, and maintaining neuronal survival in hypoxia tolerance.

## Acknowledgements

We are grateful to D. Kumar, G. Arceo-Gomez and J. Bidwell (ETSU) and M. Kirschner and L. Peshkin (Harvard) for access to laboratory equipment; to ETSU Honors college for funding (Student-Faculty collaboration grant to LYY and MCM and summer stipend for MCM), and to T. Moore for laboratory assistance.

