## Supplementary Tables and Figures for "Prior selection affects phenotypic and transcriptional response to hypoxia"

Supplementary Figures

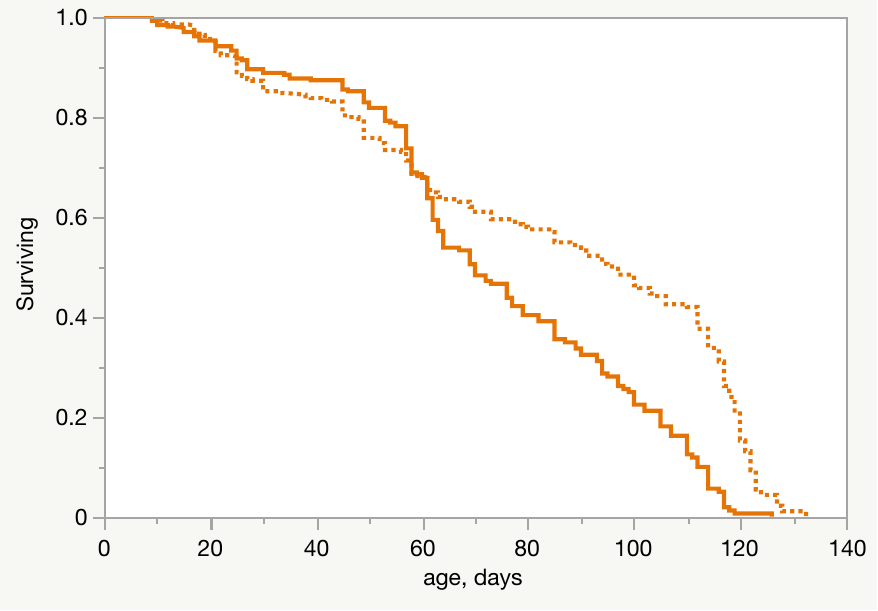

Fig. S1. Lifespan of *Daphnia* from intermittent habitats (dotted lines) and permanent habitats (solid lines) in normoxic conditions (A; green), chronic mild intermittent hypoxia (4 mg O_2_ /L twice daily; B; orange), or after a switch from 4 to 8 mg O_2_ /L at day 30. P values for Log-rank test for survival differences between groups are shown. See Table 1 for detailed survival analysis. See main text Fig.1 for the same data grouped by habitats of origin rather than hypoxia conditions.

P<0.0001

P>0.17

P>0.44

A

B

C

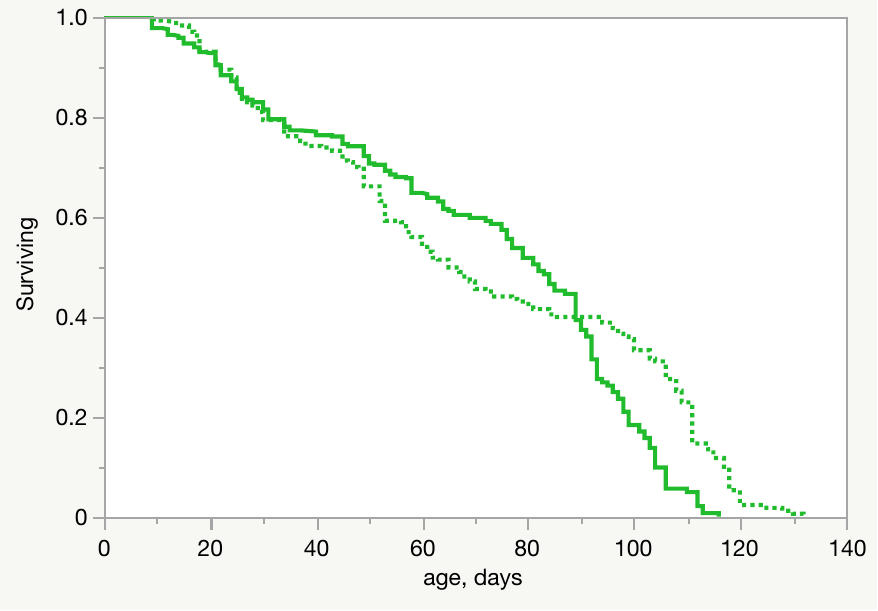

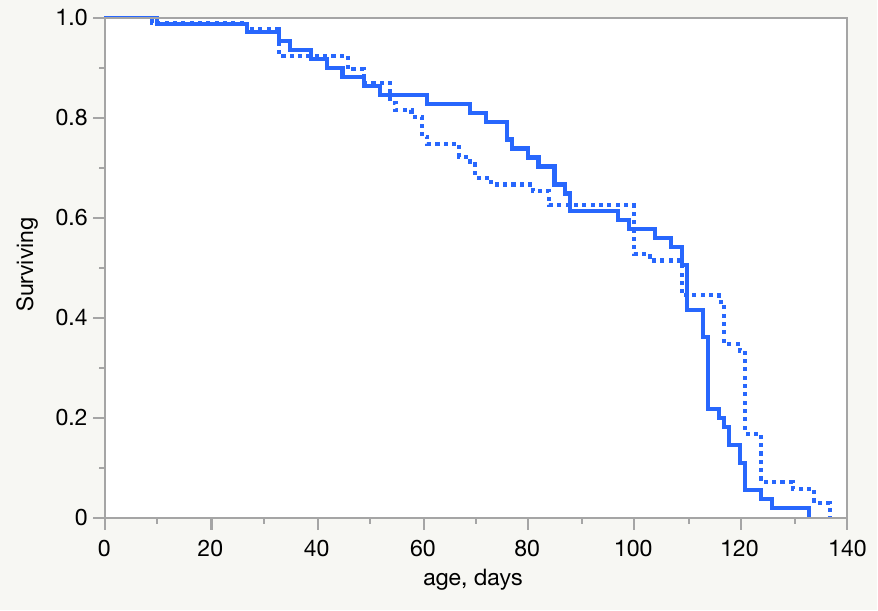

Ages: 15-20	55-60	15-20	55-60	15-20	55-60	15-20	55-60

|  | Lac, mM/mgProt |
| --- | --- |
| GB_Norm | 5.13988 |
| GB_Norm | 7.5019 |
| GB_Hyp | 5.322 |
| GB_Hyp | 9.82986 |
| IL_Norm | 4.12821 |
| IL_Norm | 5.17352 |
| IL_Hyp | 4.20966 |
| IL_Hyp | 6.12119 |

|  | Lac/Pyr |
| --- | --- |
| GB_Norm | 4.86523 |
| GB_Norm | 2.16382 |
| GB_Hyp | 6.12901 |
| GB_Hyp | 5.95504 |
| IL_Norm | 5.80523 |
| IL_Norm | 2.59707 |
| IL_Hyp | 6.81849 |
| IL_Hyp | 6.72325 |

|  | Pyr, mM/mgProt |
| --- | --- |
| GB_Norm | 1.25921 |
| GB_Norm | 3.9286 |
| GB_Hyp | 1.10916 |
| GB_Hyp | 1.88256 |
| IL_Norm | 0.812995 |
| IL_Norm | 2.55144 |
| IL_Hyp | 0.776705 |
| IL_Hyp | 1.1214 |

Fig. S2. Same data as in main text Fig. 3, for each of the two clones GB and IL separately. Whole body lactate/pyruvate ratio (A) and protein-normalized lactate and pyruvate concentrations (B,C) in young (15-20 days) and moderately aged (55-60 days) *Daphnia* reared at either normoxic control (green) or CMIH (orange) conditions. See Table 2 and Supplementary Table 2 for statistics.

A

RPKM

|  | Control | Acute Hypoxia |
| --- | --- | --- |
| intermittent | 1053.45 | 2349.28 |
| permanent | 1163.95 | 512.608 |
| intermittent | 63.7063 | 162.919 |
| permanent | 68.5559 | 39.8926 |
| intermittent | 25.459 | 64.9326 |
| permanent | 42.8485 | 12.3599 |
| intermittent | 69.3728 | 178.285 |
| permanent | 71.7107 | 42.0308 |
| intermittent | 11.8244 | 38.384 |
| permanent | 20.3222 | 12.7731 |
| intermittent | 86.8708 | 170.368 |
| permanent | 88.2335 | 38.7182 |
| intermittent | 56.5838 | 127.458 |
| permanent | 87.772 | 42.738 |
| intermittent | 346.919 | 776.862 |
| permanent | 444.567 | 271.585 |
| intermittent | 503.727 | 1090.45 |
| permanent | 537.463 | 243.327 |
| intermittent | 67.4672 | 167.225 |
| permanent | 71.0824 | 37.288 |
| intermittent | 71.1044 | 173.657 |
| permanent | 68.0016 | 37.6229 |
| intermittent | 131.727 | 347.395 |
| permanent | 60.0467 | 29.61 |
| intermittent | 58.5617 | 128.294 |
| permanent | 60.5827 | 23.6585 |
| intermittent | 30.0827 | 71.1724 |
| permanent | 33.79 | 18.2765 |

B

|  | Control | Acute Hypoxia |
| --- | --- | --- |
| intermittent | 59.1245 | 122.622 |
| permanent | 78.6042 | 42.643 |
| intermittent | 18.576 | 39.69 |
| permanent | 23.3806 | 14.8345 |
| intermittent | 103.19 | 257.213 |
| permanent | 155.617 | 89.4335 |
| intermittent | 50.5475 | 123.188 |
| permanent | 51.1043 | 31.4459 |
| intermittent | 72.0331 | 274.539 |
| permanent | 66.05 | 29.4309 |
| intermittent | 84.0364 | 241.247 |
| permanent | 189.196 | 117.367 |

Fig. S3. RPKM values of transcripts whose GOs are significantly enriched in the gene set with possible ASH x habitat type interaction. A: 14 paralogs of cuticulum proteins; B: 6 paralogs of neurotrophins. Both groups of transcripts show up-regulation in *Daphnia* from intermittent habitats and down-regulation in those from permanent habitats

RPKM

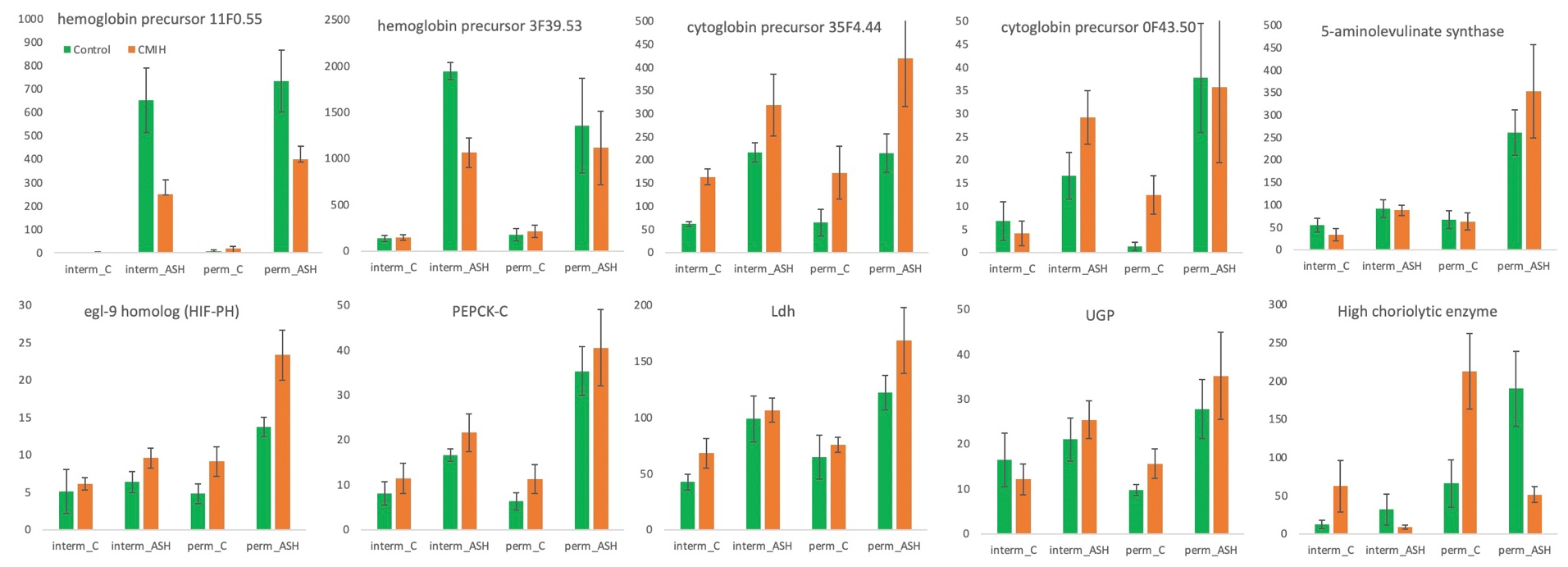

A	B	C	D	E

ASH****	ASH****	ASH***	ASH**	ASH****

CMIH****

F	G	H	I	J

ASH****	ASH****	ASH***	ASH*	CMIH***

ASH x CMIH***

Supplementary Fig. S4. Patterns of read abundance (RPKM) in select transcripts with a significant (p_adj_<0.1) effect of CMIH or ASH in the 3-way LRT analysis. *: p_adj_<0.1; **: p_adj_<0.01; ***: p_adj_<0.001; ****: p_adj_<0.0001. A-E: hemoglobins, cytoglobins and heme synthesis-related genes. F: HiF prolyl hydrolase. G – I: pyruvate metabolism and gluconeogenesis-related genes. J: choriolytic enzyme.

Supplementary tables

Table S1. Location of origin of *Daphnia* clones used. First 2 letters used in the text as clone ID.

| Clone Identifier | Location | Latitude | Longitude | Habitat type |
| --- | --- | --- | --- | --- |
| FI-FSP1-16-2 | Suur-Pellinki, Finland | 60° 10' 04" | 25° 47' 41" | Intermittent summer rock pool |
| IL-M1-8 | Jerusalem, Israel | 31° 46' 40" | 35° 13' 14" | Intermittent summer-dry pond |
| GB-EL75-69 | London, UK | 51° 31' 40" | -0° 9' 35" | Permanent pond |
| HU-K-6 | Fülöpszállás, Hungary | 46° 47' 33" | 19° 10' 54" | Permanent lake |

Table S2.

| Analysis | 3-way ANOVA | LRT: all data | LRT: by habitat of origin | LRT: subset with ASH=Control |
| --- | --- | --- | --- | --- |
| Implemented in | JMP 16 | DESeq2 | | |
| Data | RPKM | Read counts | | |
| Model | Fixed effects:  CMIH  ASH  Habitat  All interactions  Random block effect: Clones | Full model:  CMIH  ASH  Habitat  All interactions | Full model:  CMIH  ASH  Clone  All interactions | Full model:  CMIH  Habitat  CMIHxHabitat  interaction |
| Multiple tests correction | FDR | DESeq2’s padj | | |

Table S3. 2-way ANOVa of the effects of hypoxia and clones on feeding rate and body length

| Response: body length at maturity | | | | |
| --- | --- | --- | --- | --- |
| Source | DF | MS | F Ratio | Prob > F |
| hypoxia | 1 | 0.0365469 | 4.40 | 0.038 |
| clone | 3 | 2.479497 | 298.3 | <.0001 |
| hypoxia*clone | 3 | 0.0340619 | 4.10 | 0.008 |
| Error | 151 | 0.008 |  |  |
| Response: feeding rate | |  |  |  |
| Source | DF | MS | F Ratio | Prob > F |
| hypoxia | 1 | 0.331 | 12.24 | 0.0014 |
| clone | 3 | 0.155 | 5.73 | 0.003 |
| hypoxia*clone | 3 | 0.026 | 0.95 | 0.43 |
| Error | 32 | 0.027 |  |  |

| Response: respiration rate | | | | |
| --- | --- | --- | --- | --- |
| Source | DF | MS | F Ratio | Prob > F |
| AssayO2 | 1 | 22.90 | 2.047 | 0.16 |
| hypoxia | 1 | 0.12 | 0.011 | 0.92 |
| AssayO2*hypoxia | 1 | 0.12 | 0.010 | 0.92 |
| clone | 3 | 2.72 | 0.243 | 0.87 |
| AssayO2*Ccone | 3 | 0.84 | 0.075 | 0.97 |
| hypoxia*clone | 3 | 2.63 | 0.236 | 0.87 |
| AssayO2* hypoxia *clone | 3 | 12.29 | 1.099 | 0.35 |
| Error | 101 | 11.18 |  |  |

| Response: respiration rate, measured at acclimation O_2_ concentration | | | | |
| --- | --- | --- | --- | --- |
| Source | DF | MS | F Ratio | Prob > F |
| hypoxia | 1 | 19.78 | 2.29 | 0.13 |
| clone | 3 | 7.21 | 0.84 | 0.48 |
| hypoxia *clone | 3 | 1.75 | 0.20 | 0.89 |
| Error | 70 | 8.63 |  |  |

| Response: mean clutch size at age 18 days | | | |  |  |
| --- | --- | --- | --- | --- | --- |
| Source | DF | MS | F Ratio | | Prob > F |
| hypoxia | 1 | 0.22 | 0.0276 | | 0.87 |
| clone | 3 | 11.20 | 1.3898 | | 0.27 |
| hypoxia *clone | 3 | 2.00 | 0.2482 | | 0.86 |
| Error | 24 | 8.06 |  | |  |

Table S3 continued

| Response: mean clutch size at age 38 days | | | |  |  |
| --- | --- | --- | --- | --- | --- |
| Source | DF | MS | F Ratio | | Prob > F |
| hypoxia | 1 | 0.84 | 1.86 | | 0.21 |
| clone | 3 | 2.95 | 6.54 | | 0.015 |
| hypoxia *clone | 3 | 0.62 | 1.37 | | 0.32 |
| Error | 8 | 0.45 |  | |  |

| Response: mean clutch size at age 66 days | | | |  |  |
| --- | --- | --- | --- | --- | --- |
| Source | DF | MS | F Ratio | | Prob > F |
| hypoxia | 1 | 242.67 | 32.1 | | 0.0013 |
| clone | 3 | 7.98 | 1.06 | | 0.43 |
| hypoxia *clone | 3 | 27.61 | 3.65 | | 0.08 |
| Error | 6 | 7.56 |  | |  |

Table S4. Three-way ANOVA of the effects of CMIH, age, and clones on protein content-normalized lactate and pyruvate concentrations (Fig. 3 and S2). “Plate” is a random block effect.

Response Pyr, mM/mgProt

| Source | DF | MS | F Ratio | Prob > F |
| --- | --- | --- | --- | --- |
| CMIH | 1 | 49.36 | 4.82 | 0.03 |
| clone | 1 | 90.05 | 8.80 | 0.0037 |
| CMIH*clone | 1 | 8.99 | 0.88 | 0.35 |
| age | 1 | 0.022 | 0.002 | 0.96 |
| CMIH *age | 1 | 40.45 | 3.95 | 0.049 |
| clone*age | 1 | 15.90 | 1.55 | 0.21 |
| CMIH *clone*age | 1 | 7.49 | 0.73 | 0.39 |
| plate | 1 | 109.65 | 10.72 | 0.0014 |
| Error | 103 | 10.23 |  |  |

Response Lac, mM/mgProt

| Source | DF | MS | F Ratio | Prob > F |
| --- | --- | --- | --- | --- |
| CMIH | 1 | 23.51 | 20.97 | <.0001 |
| clone | 1 | 15.0 | 13.38 | 0.0004 |
| CMIH*clone | 1 | 1.08 | 0.96 | 0.33 |
| age | 1 | 28.23 | 25.18 | <.0001 |
| CMIH *age | 1 | 19.17 | 17.10 | <.0001 |
| clone*age | 1 | 3.44 | 3.072 | 0.083 |
| CMIH *clone*age | 1 | 0.56 | 0.50 | 0.48 |
| plate | 1 | 0.85 | 0.75 | 0.38 |
| Error | 103 | 1.12 |  |  |

Supplementary Table S5. Mixed model 2-way ANOVa of the coordinates in the first 3 principal components in the space of 578 transcripts with at least 1 significant non-adjusted p-value. CMIH and ASH as fixed effects, clones as a random block. Analysis for clones from: intermittent and permanent habitats of origin separately.

| Habitat of origin: | | | | intermittent | | | permanent | |
| --- | --- | --- | --- | --- | --- | --- | --- | --- |
|  | Source | df | df_Den_ | F Ratio | Prob > F | F Ratio | | Prob > F |
| Response: PC1 | CMIH | 1 | 19 | 2.3416 | 0.14 | 0.4789 | | 0.50 |
|  | ASH | 1 | 19 | 16.226 | **0.0007** | 1.877 | | 0.18 |
|  | CMIH*ASH | 1 | 19 | 0.0562 | 0.82 | 0.0677 | | 0.80 |
| Response: PC2 | CMIH | 1 | 19 | 0.1161 | 0.74 | 0.0519 | | 0.82 |
|  | ASH | 1 | 19 | 18.6566 | **0.0004** | 0.7377 | | 0.40 |
|  | CMIH*ASH | 1 | 19 | 0.0091 | 0.93 | 0.0015 | | 0.97 |
| Response: PC3 | CMIH | 1 | 19 | 6.6618 | *0.0183* | 3.4484 | | 0.079 |
|  | ASH | 1 | 19 | 40.3107 | **<.0001** | 34.7833 | | **<.0001** |
|  | CMIH*ASH | 1 | 19 | 0.0036 | 0.95 | 0.0031 | | 0.96 |
